# Development of in vitro test methods in a model of human pancreatic β-cells to identify metabolism disrupting chemicals with diabetogenic activity

**DOI:** 10.1101/2022.03.22.485271

**Authors:** Reinaldo Sousa Dos Santos, Regla María Medina-Gali, Ignacio Babiloni-Chust, Laura Marroqui, Angel Nadal

## Abstract

There is a need to develop identification tests for Metabolism Disrupting Chemicals (MDCs) with diabetogenic activity. Here we used the human EndoC-βH1 β-cell line, the rat β-cell line INS-1E and dispersed mouse islet cells to assess the effects of endocrine disruptors on cell viability and glucose-stimulated insulin secretion (GSIS). We tested six chemicals at concentrations within human exposure (from 0.1 pM to 1 μM). Bisphenol-A (BPA) and tributyltin (TBT) were used as controls while four other chemicals, namely perfluorooctanoic acid (PFOA), triphenylphosphate (TPP), triclosan (TCS) and dichlorodiphenyldichloroethylene (DDE), were used as “unknowns”. Regarding cell viability, BPA and TBT increased cell death as previously observed. Their mode of action involved the activation of estrogen receptors and PPARγ, respectively. ROS production was a consistent key event in BPA- and TBT-treated cells. None of the other MDCs tested modified viability or ROS production. Concerning GSIS, TBT increased insulin secretion while BPA produced no effects. PFOA decreased GSIS, suggesting that this chemical could be a “new” diabetogenic agent. Our results indicate that the EndoC-βH1 cell line is a suitable human β-cell model for testing diabetogenic MDCs. Optimization of the test methods proposed here could be incorporated into tier protocols for the identification of MDCs.

## 1. Introduction

Diabetes prevalence has been continuously growing in recent decades, reaching pandemic proportions [1]. Genetic and environmental factors play a role in diabetes etiology. While the genetic background may predispose individuals to the disease, environmental factors, including exposure to chemical pollutants that interfere with the endocrine system (also known as endocrine-disrupting chemicals or EDCs), may act as triggers to diabetes development. Type 1 diabetes is an autoimmune disease characterized by pancreatic β-cell apoptosis as the result of an immune attack [2,3]. Although it is still unclear whether there is a link between type 1 diabetes and exposure to EDCs, some evidence indicate that EDC exposure, especially during development, may play a role in the pathogenesis of type 1 diabetes [4]. Type 2 diabetes is the result of β-cell dysfunction and eventually death, which usually happen in a background of insulin resistance. β-cell dysfunction represents an early phenomenon in the development of type 2 diabetes as it occurs before dysglycemia advances [5]. In some special cases, insulin hypersecretion may also lead to dysglycemia [5,6]. After two decades of research, the relationship between exposure to EDCs and type 2 diabetes development has grown sufficiently strong to consider these environmental pollutants as a potential cause of type 2 diabetes [7–9]. Metabolism-disrupting chemicals (MDCs) are defined as endocrine disruptors that alter susceptibility to metabolic disorders, including diabetes, obesity, and non-alcoholic fatty liver disease [8,10]. Despite growing evidence suggesting a relationship between exposure to MDCs and susceptibility to metabolic diseases in cellular and animal models as well an in epidemiological studies, there is a lack of test methods to identify potential MDCs [11]. GOLIATH is a project within the Horizon 2020 program of the European Commission whose aim is to address the urgent need to develop test protocols that allow the identification of MDCs that cause relevant toxic effects through an endocrine mode of action [11]. One of our objectives within the GOLIATH project is to design test methods to identify MDCs that may pose a risk for the development of diabetes.

As the insulin produced and secreted by β-cells is the only hormone in our body that lowers blood glucose levels, any disruption in its secretion (*e.g*., due to β-cell malfunction and/or death) poses a serious risk for diabetes onset. Therefore, it is essential to develop robust tests to detect MDCs that might decrease β-cell viability and/or disrupt its function. In the present work we sought to develop in vitro test methods that could allow the identification of metabolism disrupting activity in β-cells. We used two β-cell lines, namely the rat insulinoma-derived INS-1E [12] and the human EndoC-βH1 [13]. INS-1E cells are a stable β-cell line surrogate with less heterogeneity than the non-clonal INS-1 cell line from which INS-1E were cloned [12]. EndoC-βH1 cells present similar metabolic function to INS-1 832/13 and similar insulin secretion in response to glucose to that observed in human islets; this means that EndoC-βH1 cells are as useful as INS-1 cells but with the advantage of being from human origin [14]. Both cell lines were exposed to six MDCs, namely Bisphenol A (BPA), Tributyltin (TBT), Perfluorooctanoic acid (PFOA), Triphenylphosphate (TPP), Triclosan (TCS), and Dichlorodiphenyldichloroethylene (DDE). The concentration range tested herein was based on biomonitoring studies for BPA [15], TBT [16], PFOA [17], TPP [18], TCS [19] and DDE [20]. BPA and TBT have been previously tested in β-cells. BPA and TBT decreased β-cell viability in different β-cell models, namely INS-1E cells [21] and RINm5F cells [22]. Moreover, both chemicals have been characterized as MDCs that alter insulin secretion in primary mouse islets/β-cells [22–24], rat islets and insulinomas [25–27], and human islets [28,29]. Therefore, we used these two chemicals as positive controls to evaluate our human β-cell model. Despite some evidence of their potential as MDCs, the other four chemicals, PFOA, TPP, TCS and DDE, were considered “unknown” chemicals due to the lack of data about their effects on β-cells. To test these chemicals, we followed an adverse outcome pathway framework, in which we studied the molecular initiated event (MIE) by pharmacology and two key events, namely gene expression and reactive oxygen species (ROS) production. Finally, we evaluated β-cell survival and glucose-stimulated insulin secretion (GSIS) as adverse effects.

## 2. Results

### 2.2. BPA and TBT induce β-cell death

To assess whether different MDCs could affect cell survival, we measured cell viability by two different methods, namely the MTT assay and staining with the DNA-binding dyes Hoechst 33342 and propidium iodide (HO/PI). The former is a colorimetric method that indicates cell viability based on metabolic activity [30,31], while the latter is a fluorescent method that allows discrimination between viable, apoptotic, and necrotic cells [32]. We exposed EndoC-βH1 and INS-1E cells to a range of concentrations of each MDC and cell viability was measured by MTT assay 48 h and 72 h following treatment (Figure 1, Supplementary Figure 1, and data not shown). Of the six chemicals tested, only BPA (Figure 1A, Supplementary Figure 1A) and TBT (Figure 1B, Supplementary Figure 1B) decreased cell viability, both acting in a dose-dependent manner. When compared to their respective vehicles, the highest BPA dose (i.e., 1 μM) induced a 15-20% decrease, whereas the highest TBT dose (i.e., 200 nM) decreased viability by 24-27% after exposure for 48 h, depending on the cell model used.

**Figure 1.**
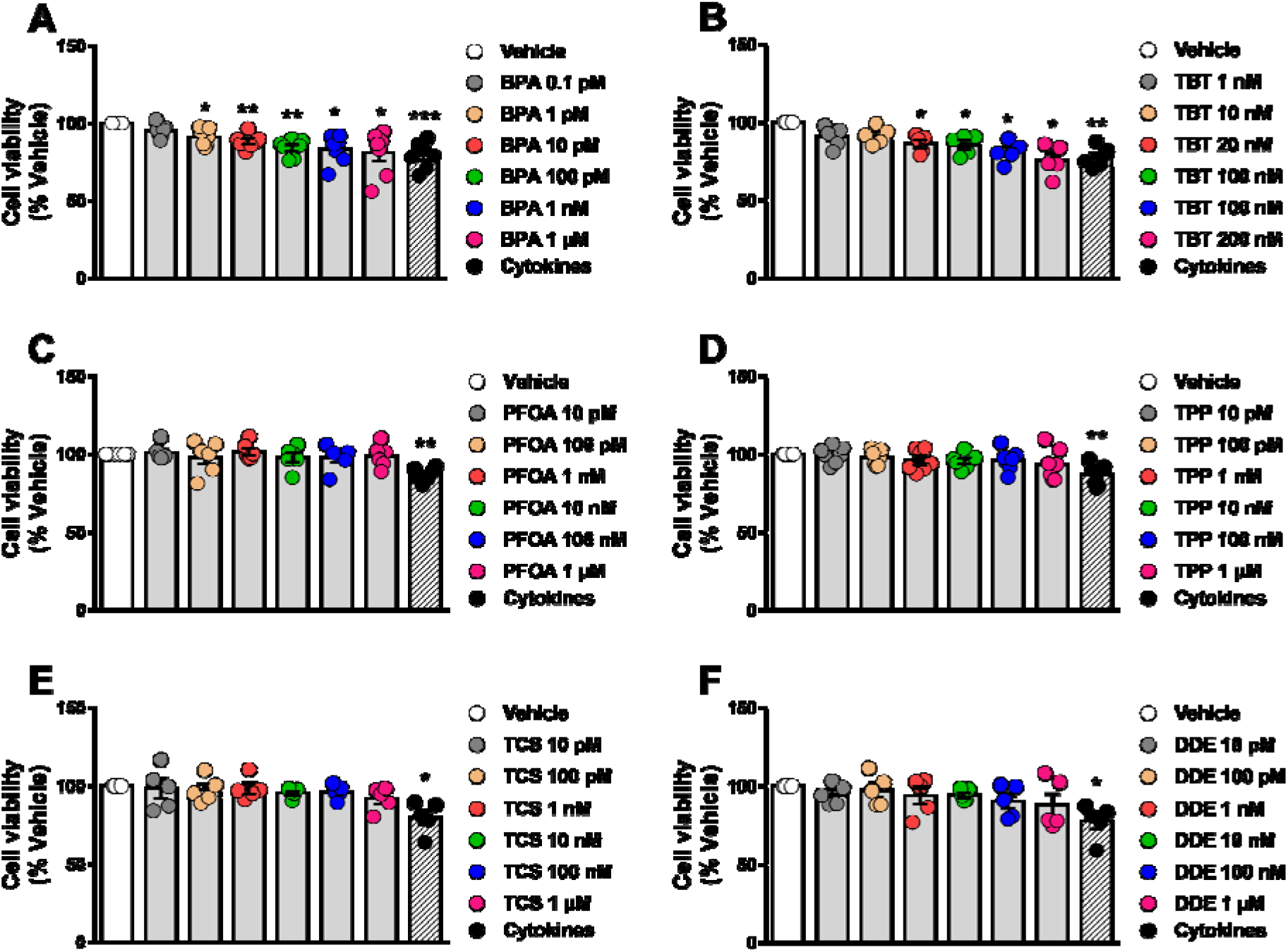
β-cell viability upon MDC exposure. EndoC-βH1 cells were treated with vehicle (DMSO) or different doses of BPA (A), TBT (B), PFOA (C), TPP (D), TCS (E), or DDE (F) for 48 h. A cocktail of the cytokines IL-1β + IFNγ (50 and 1000 U/ml, respectively) was used as a positive control. Cell viability was evaluated by MTT assay. Results are expressed as % vehicle-treated cells. Data are shown as means ± SEM of five to seven independent experiments. *p≤0.05, **p≤0.01 and ***p≤0.001 vs Vehicle. MDCs vs Vehicle by one-way ANOVA; Cytokines vs Vehicle by two-tailed Student’s t test.

As seen by the MTT assay, assessment of β-cell viability by HO/PI showed that BPA and TBT induced apoptosis in EndoC-βH1 and INS-1E cells upon 24 h treatment (Figure 2, Supplementary Figure 2). Due to the sensitivity of this method, we observed that BPA doses within the picomolar range increased apoptosis in both cell lines (Figure 2A,D). To compare these results with a more physiological cell system, we used dispersed mouse islet cells. As observed in both β-cell lines, BPA and TBT also induced apoptosis in dispersed islets (Figure 2G,H), whereas PFOA, TPP, TCS, and DDE did not affect the viability of these cells (Figure 2I). Depending on the cell model, 1 μM BPA induced 1.7 to 2.5-fold increase in apoptosis, while 200 nM TBT promoted a 1.5 to 3-fold increase in apoptosis. These findings indicate that MDCs affect the viability of the human EndoC-βH1 cells, the rat line INS-1E and the primary mouse β-cells in a very similar way. Based on the findings described above, we used only EndoC-βH1 cells to confirm our results with a third method, namely caspase 3/7 activity assay. We observed that caspase 3/7 activity was augmented by BPA and TBT exposure but no by exposure to PFOA, TPP, TCS, and DDE (Supplementary Figure 3).

**Figure 2.**
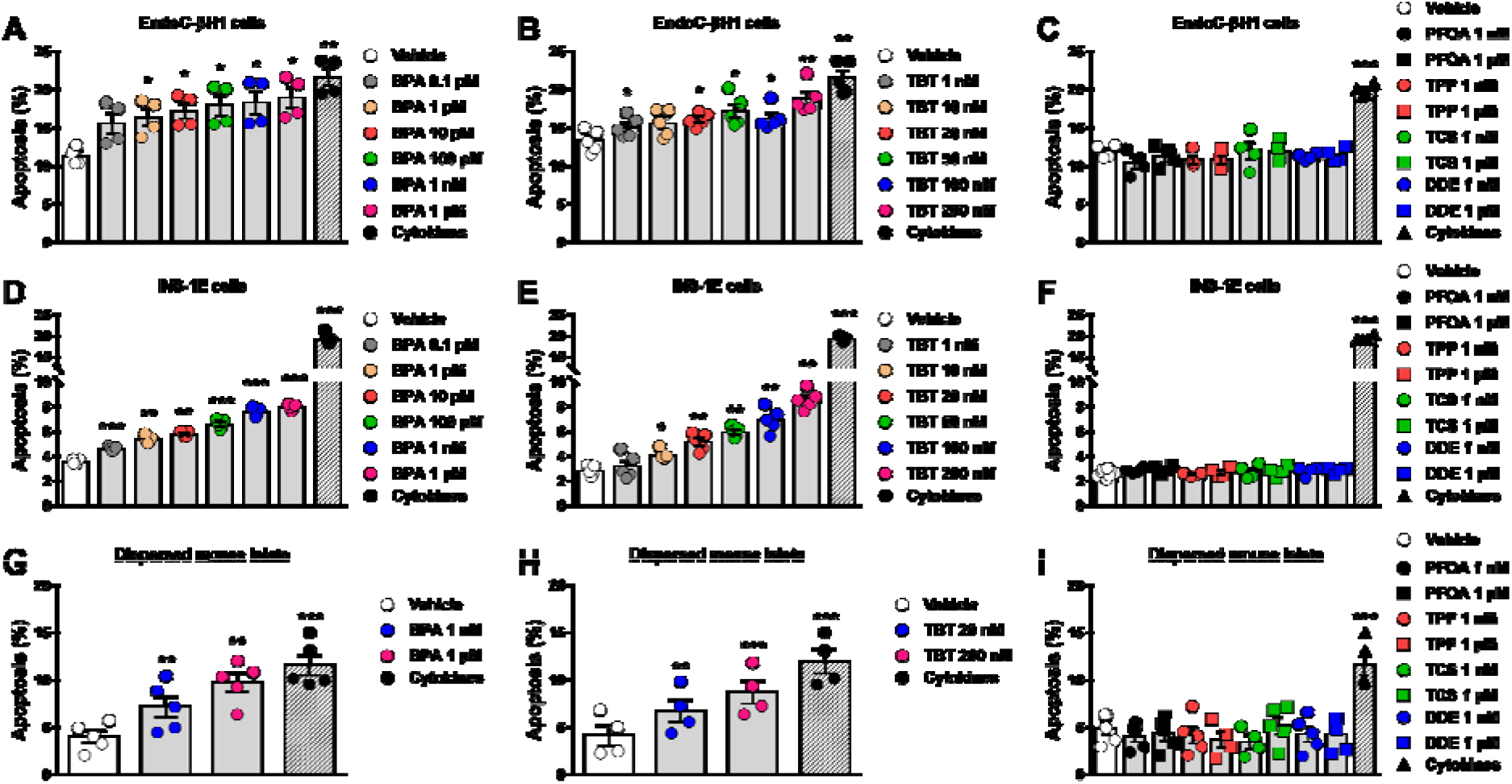
β-cell apoptosis upon MDC exposure. EndoC-βH1 (A–C) and INS-1E cells (D–F), or dispersed mouse islets (G–I) were treated with vehicle (DMSO) or different doses of BPA (A, D, and G), TBT (B, E, and H), PFOA, TPP, TCS, or DDE (C, F, and I) for 24 h (A–F) or 48 h (G–I). A cocktail of the cytokines IL-1β + IFNγ (10 and 100 U/ml, respectively for INS-1E cells; 50 and 1000 U/ml, respectively, for EndoC-βH1 cells and dispersed mouse islets) was used as a positive control. Apoptosis was evaluated using HO and PI staining. Data are shown as means ± SEM of four to five independent experiments. *p≤0.05, **p≤0.01 and ***p≤0.001 vs Vehicle. MDCs vs Vehicle by one-way ANOVA; Cytokines vs Vehicle by two-tailed Student’s t test.

A cocktail of proinflammatory cytokines, namely interleukin-1β (IL-1β) and interferon-γ (IFNγ), was used as positive control. As expected, this mix of cytokines decreased viability, induced apoptosis or increased caspase 3/7 activity in all models tested (Figure 1,2, Supplementary Figure 1,3).

### 2.2. Estrogen receptors are involved in BPA-induced β-cell apoptosis

In β-cells, BPA effects on insulin content and secretion as well as in apoptosis involve the activation of estrogen receptors ERα and ERβ [23,28,33]. Figures 3A and B show that the pure estrogen receptors antagonist ICI 182,780 completely blocked BPA-induced apoptosis in EndoC-βH1 (Figure 3A) and INS-1E cells (Figure 3B). These results indicate that the MIE responsible for the BPA-induced β-cell death described in Figures 1 and 2 involves estrogen receptors.

**Figure 3.**
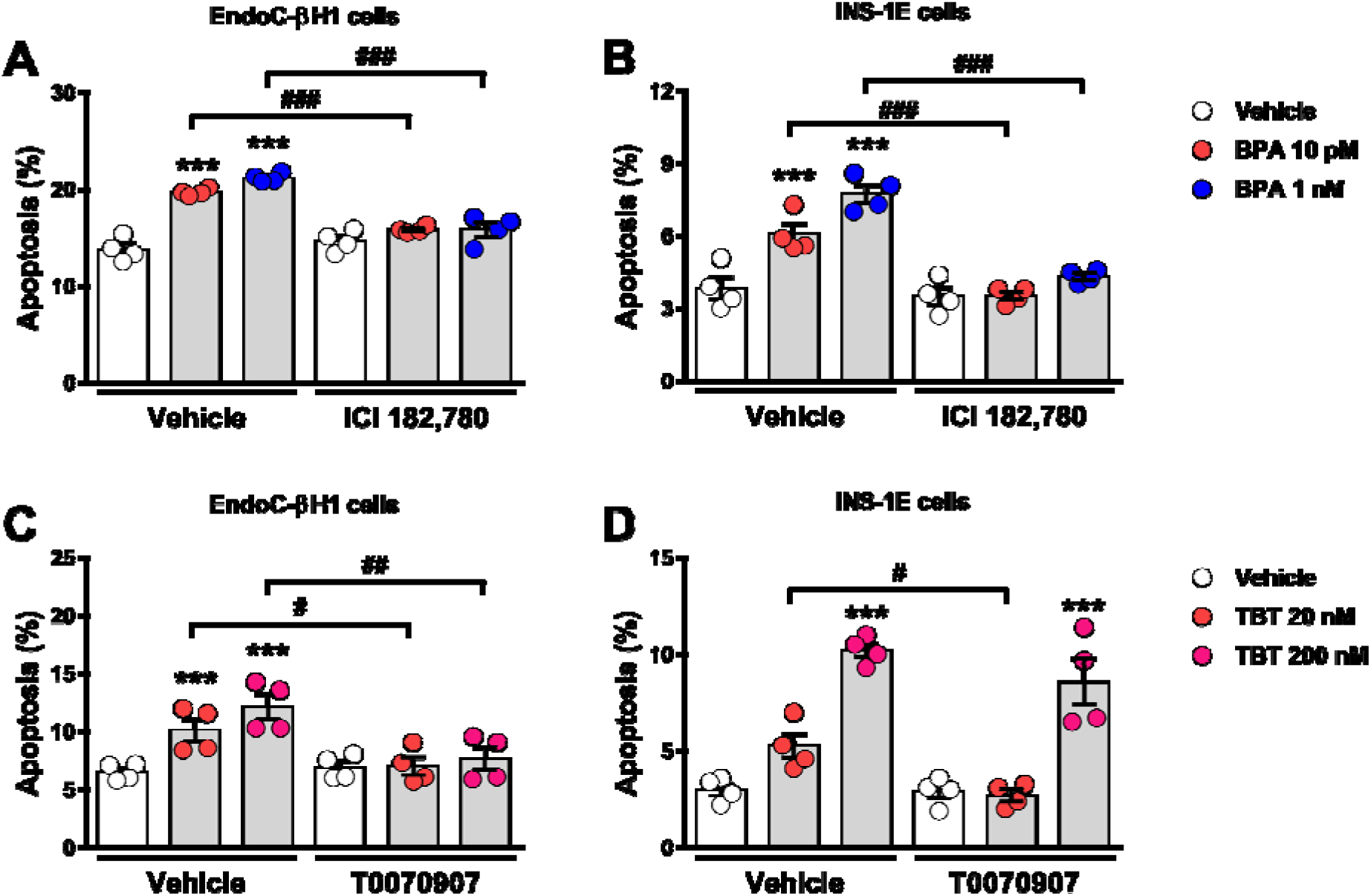
Estrogen receptors and PPARγ are involved in BPA- and TBT-induced β-cell apoptosis, respectively. EndoC-βH1 (A) and INS-1E (B) cells were treated with vehicle (DMSO) or BPA (10 pM or 1 nM) in the absence or presence of 1 μM ICI 182,780 for 24 h. EndoC-βH1 (C) and INS-1E (D) cells were treated with vehicle (DMSO) or TBT (20 nM or 200 nM) in the absence or presence of 100 nM T0070907 for 24 h. Apoptosis was evaluated using HO and PI staining. Data are shown as means ± SEM of four independent experiments. ***p≤0.001 vs its respective Vehicle; #p≤0.05, ##p≤0.01, and ###p≤0.001 as indicated by bars. Two-way ANOVA.

### 2.3. PPARγ is involved in TBT-induced β-cell apoptosis

TBT acts as an agonist of both the peroxisome proliferator-activated receptor γ (PPARγ) and retinoid X receptors (RXRs) [34,35]. To assess whether PPARγ activation was part of the MIE whereby TBT induces apoptosis, we used the PPARγ antagonist T0070907. In EndoC-βH1 cells, T0070907 blocked TBT-induced apoptosis at both doses (Figure 3C), while in INS-1E cells, T0070907 prevented the apoptotic effect of TBT at a low (20 nM) but not at a high dose (200 nM) (Figure 3D).

Rosiglitazone, a well-known PPARγ-selective activator, not only did not induce apoptosis, but it also abrogated TBT-induced apoptosis in EndoC-βH1 and INS-1E cells (Supplementary Figure 4). These findings suggest that PPARγ activation is part of the MIE whereby TBT induces β-cell apoptosis and show that both PPARγ agonists, rosiglitazone and TBT, have opposite effects on β-cell viability.

### 2.4. BPA and TBT, but not PFOA, induce ROS generation

Previous work demonstrated that both BPA and TBT induced oxidative stress in β-cells [21,25]. In line with these findings, we observed that BPA and TBT induced ROS generation in both cell lines explored here (Figure 4A-D). PFOA, which did not induce apoptosis at any of the concentrations used, failed to produce ROS at both low (1 nM) and high (1 μM) doses (Figure 4 E,F). Menadione, which is known for inducing ROS production, was used as a positive control. These results indicate that ROS production is a key event involved in BPA- and TBT-induced β-cell apoptosis.

**Figure 4.**
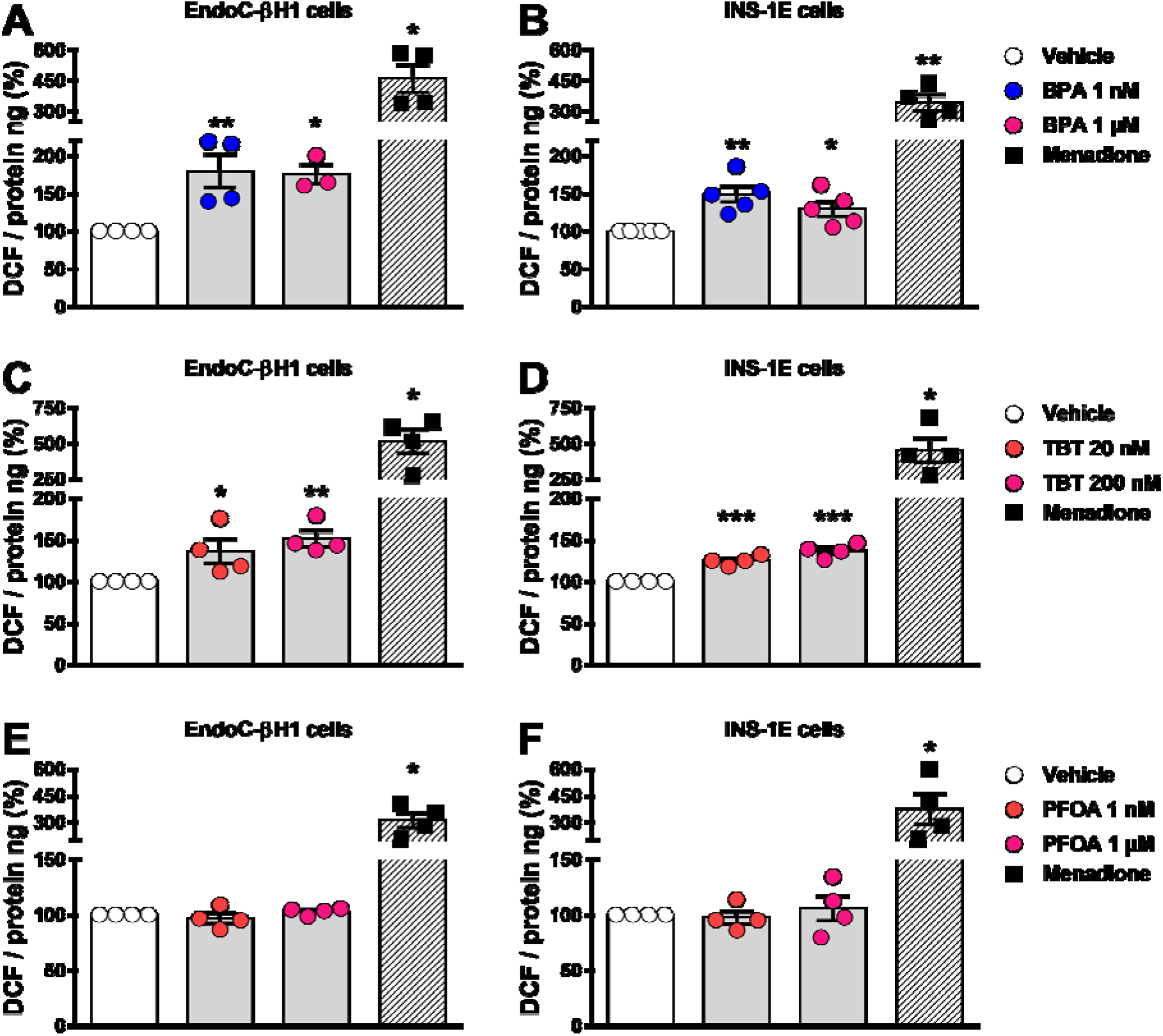
ROS production upon MDC exposure. EndoC-βH1 (A, C and E), and INS-1E (B, D and F) cells were treated with vehicle (DMSO) or different doses of BPA (A and B), TBT (C and D), or PFOA (E and F) for 24 h. Menadione (15 μM for 90 minutes) was used as a positive control. ROS production was measured by oxidation of the fluorescent probe 2’,7’-dichlorofluorescein diacetate (DCF) and normalized by total protein. Results are expressed as % vehicle-treated cells. Data are shown as means ± SEM of three to five independent experiments. *p≤0.05, **p≤0.01 and ***p≤0.001 vs Vehicle. MDCs vs Vehicle by one-way ANOVA; Menadione vs Vehicle by two-tailed Student’s t test.

### 2.5. β-cell function is disturbed by different MDCs

Our next step was to explore whether the six MDCs tested herein could perturb β-cell function. For this purpose, we measured glucose-stimulated insulin secretion (GSIS) and insulin content in EndoC-βH1 cells upon exposure to different doses of each MDC for 48 h (Figure 5). BPA (Figure 5A) and TCS (Figure 5E) did not modify GSIS, despite a trend observed in cells treated with TCS 1 μM. An increase in insulin secretion was observed upon exposure to the highest doses of TBT (200 nM; Figure 5B), TPP (1 μM; Figure 5D), and DDE (1 μM; Figure 5F). Curiously, we also observed an increase in insulin secretion at low glucose in EndoC-βH1 cells treated with 100 pM TPP (Figure 5D). PFOA promoted the most changes in β-cell function as exposure to several doses of this MDC decreased insulin secretion both at low and high glucose concentrations (Figure 5C). Interestingly, we found that PFOA modulated insulin secretion in a non-monotonic dose response-dependent manner, in which exposure from 10 pM to 100 nM PFOA reduced insulin secretion, while exposure to 1 μM did not significantly changed secretion (Figure 5C).

**Figure 5.**
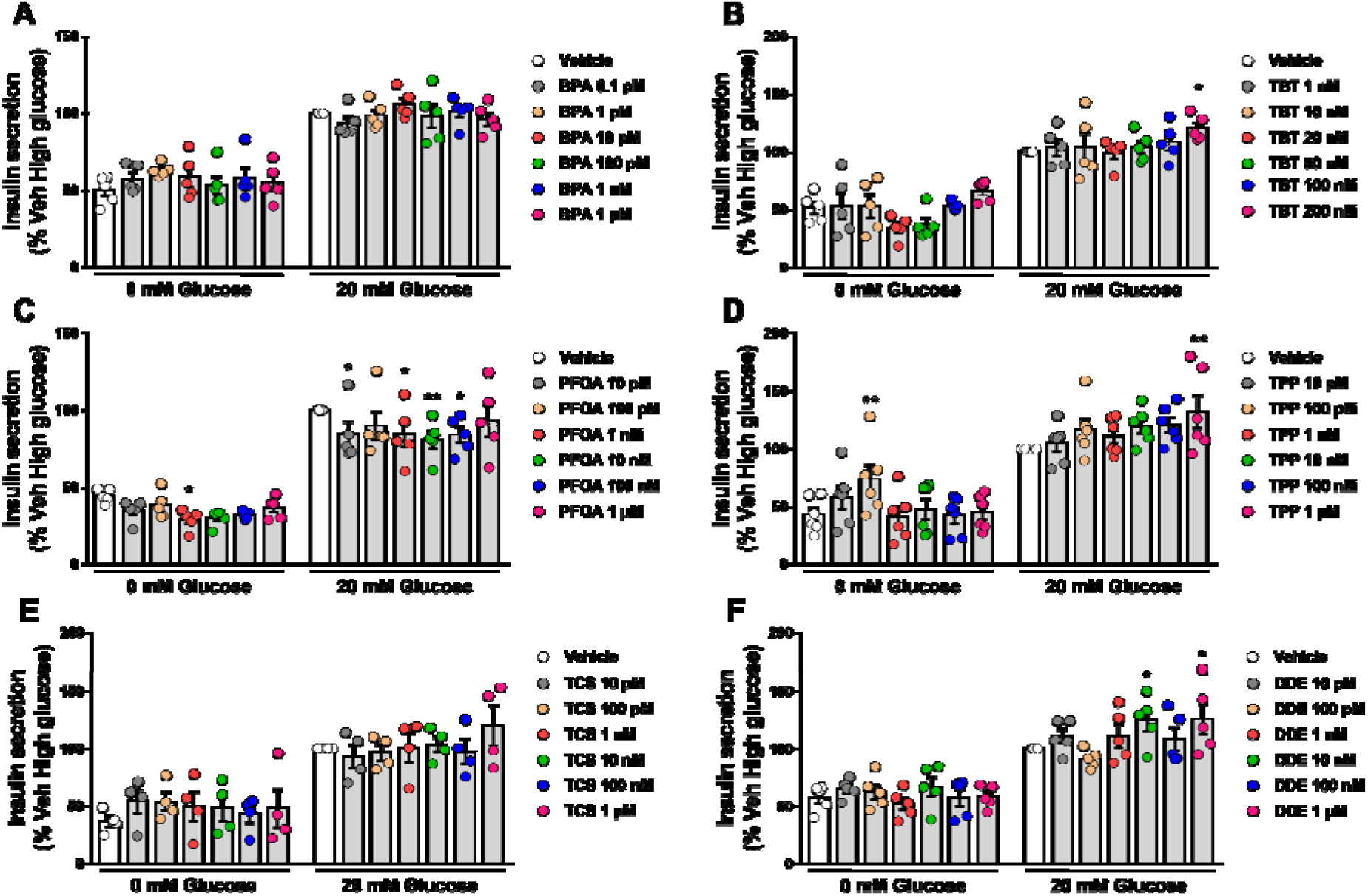
Glucose-stimulated insulin secretion upon MDC exposure. EndoC-βH1 cells were treated with vehicle (DMSO) or different doses of BPA (A), TBT (B), PFOA (C), TPP (D), TCS (E), or DDE (D) for 48 h. Insulin secretion was measured at 0 and 20 mM glucose, and insulin released into the medium was measured by ELISA. Data are normalized to insulin secretion at high glucose (20 mM) in vehicle-treated cells (considered as 100%). Data are shown as means ± SEM of four to six independent experiments. *p≤0.05, **p≤0.01 and ***p≤0.001 vs its respective Vehicle. Two-way ANOVA.

Exposure to BPA, PFOA, TPP, TCS, and DDE did not modify insulin content (Supplementary Figure 5); on the contrary, TBT promoted a slight increase at 20 nM and a 20% decrease in insulin content at 200 nM (Supplementary Figure 5).

### 2.6. Effects of MDCs on β-cell gene expression

Finally, we examined whether the different MDCs could modulate the expression of genes involved in β-cell identity (*PDX1* and *MAFA*) and β-cell function (*INS, GLUT2* and *GCK*). After 24 h exposure, mRNA expression was measured in EndoC-βH1 and INS-1E cells. TBT significantly decreased *MAFA* expression, reaching about 50% inhibition in EndoC-βH1 and INS-1E cells (Figure 6B, Supplementary Figure 6B). In EndoC-βH1 cells, TBT also induced a trend towards downregulation of *PDX1*, but these results did not reach statistical significance (Figure 6E). Of note, *Pdx1* expression decreased upon exposure to TBT 200 nM in INS-1E cells (Supplementary Figure 6E). In addition, TBT increased *GLUT2* expression in EndoC-βH1 cells but not in INS-1E cells (Figure 6K, Supplementary Figure 6N). No major significant changes in gene expression were observed when cell lines were treated with BPA, PFOA, TPP, TCS and DDE (Figure 6, Supplementary Figure 6).

**Figure 6.**
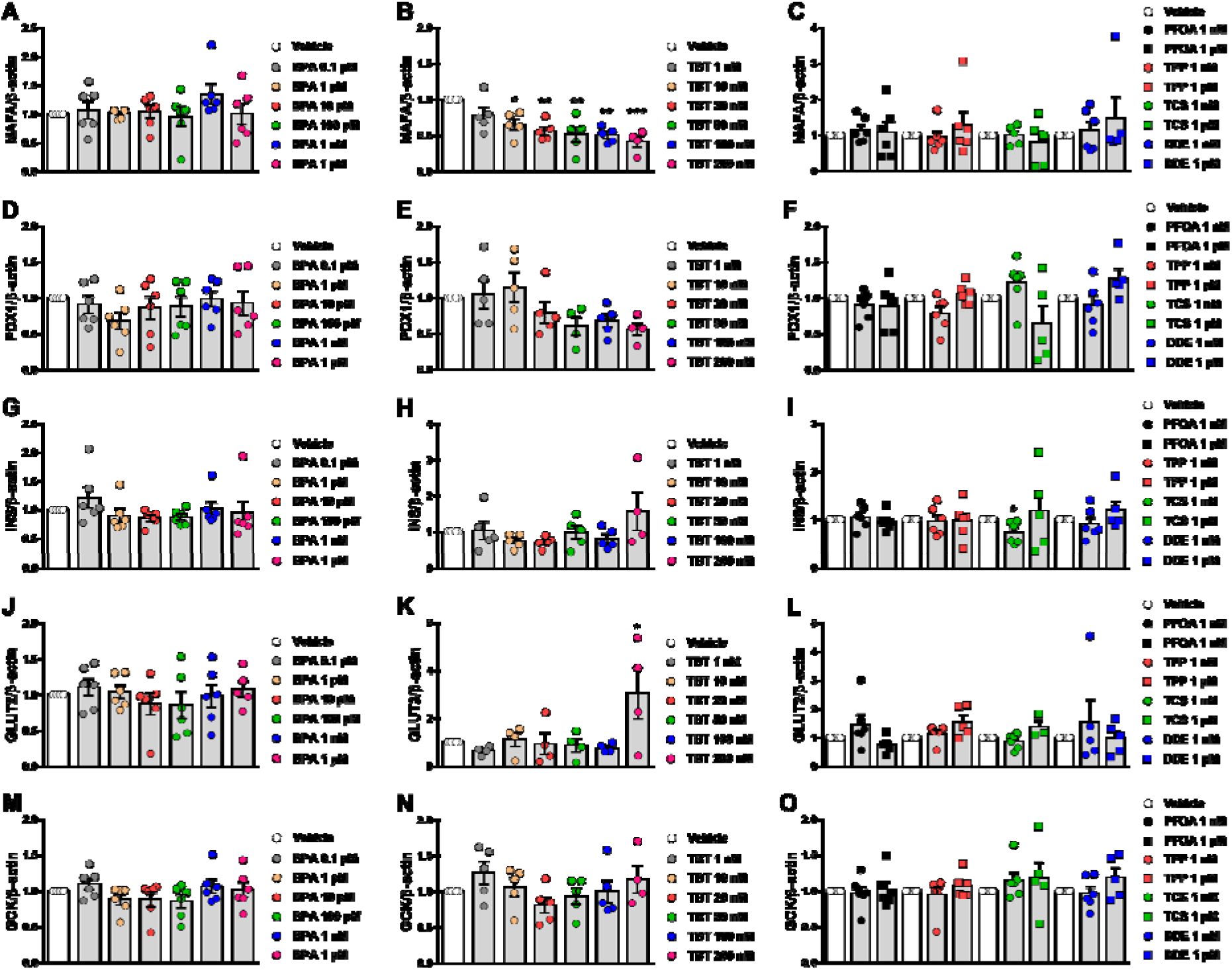
Gene expression upon MDC exposure in EndoC-βH1 cells. mRNA expression of *MAFA* (A-C), *PDX1* (D-F), *INS* (G-I), *GLUT2* (J-L), and *GCK* (M-O) was measured in EndoC-βH1 cells treated with vehicle (DMSO) or different doses of BPA (A, D, G, J and M), TBT (B, E, H, K and N), PFOA, TPP, TCS, or DDE (C, F, I, L and O) for 24 h. mRNA expression was measured by qRT-PCR and normalized to the housekeeping gene β-actin, and then by vehicle-treated cells (considered as 1). Data are shown as means ± SEM of four to six independent experiments. *p≤0.05, **p≤0.01 and ***p≤0.001 vs its respective Vehicle. One-way ANOVA.

## 3. Discussion

Diabetes develops because there is a decrease in functional pancreatic β-cell mass, which may occur after disruption of β-cell secretory capacity, β-cell death, or both. Therefore, it is plausible to say that GSIS and β-cell viability are the two main deterministic endpoints to study diabetogenic factors.

Our main objective in this work was to test and validate whether the EndoC-βH1 cell line would be an effective human β-cell model for developing test methods to assess β-cell viability and function. The development of these tests in a human β-cell model will help to identify contaminants with the potential to cause diabetes through an endocrine mode of action. The human EndoC-βH1 cell line has been used worldwide as a useful tool to increase our understanding of human islet biology and diabetes [36,37]. These cells recapitulate features of adult primary β-cells, and their open chromatin, transcriptomics and proteome landscapes were reported to be very similar to adult human β-cells [38,39]. In fact, EndoC-βH1 cells express all genes and proteins that lead to the phenotype of a typical β-cell [40], including a similar set of ion channels, electrical activity, and calcium signaling in response to glucose as compared to primary human β-cells [41]. Moreover, these cells are an appropriate human β-cell model for novel drug screening [42]. For the experimental protocols used in the present work we followed the recommendations of Univercell-Biosolutions, the company that supplies the EndoC-βH1 cells, including all culture media. This cell culture protocol has been previously used in key studies validating EndoC-βH1 cells as a suitable model of human β-cells [40–44].

As previous works have shown that BPA and TBT affect β-cell viability and function in rodent β-cell lines and mouse β-cells, we compared our results in EndoC-βH1 cells with those obtained in the rat INS-1E cell line and in dispersed cells isolated from mouse islets.

### 3.1. β-cell viability tests

Our results showed that BPA and TBT induced β-cell death, whereas PFOA, TPP, TCS and DDE did not modify cell survival at the concentrations tested. Similar results were obtained using two different approaches to measure β-cell viability, namely the MTT assay and HO/PI DNA-binding dyes, in both cell lines studied. We confirmed our HO/PI findings in dispersed mouse islet cells. Moreover, we used a third method, *i.e*., caspase 3/7 activity assay, to confirm our data obtained in EndoC-βH1 cells. A cocktail of proinflammatory cytokines (IL-1β + IFNγ) normally employed in β-cell research [45,46] was used as a positive control to assure the methodology was working.

There is plenty of evidence showing that BPA is an MDC that alter metabolism through an endocrine mode of action [7,47]. At nanomolar concentrations, BPA increased apoptosis following mitochondrial dysfunction in INS-1 cells [21,25]. Pancreatic β-cells from rodent pups perinatally treated with low doses of BPA (50 μg/kg/day, oral exposure) exhibited hypertrophy mitochondria and rough endoplasmic reticulum compared to control pups [48]. Male mouse pups prenatally treated with 10 μg/kg/day BPA showed lower pancreatic β-cell mass associated with increased caspase 3 activity and altered gene expression associated with mitochondrial function [49].

Our present findings regarding the effect of BPA on cell viability in EndoC-βH1, INS-1E and dispersed mouse islet cells mirrored previous data in β-cell models. In agreement with our previous data describing a crosstalk between the estrogen receptors G protein-coupled estrogen receptor 1, ERα and ERβ [33], BPA-induced apoptosis at low doses was abolished by the pure ERα and ERβ antagonist ICI 182,780. Thus, the MIE in the action of BPA involves estrogen receptors, including the G protein-coupled estrogen receptor 1, which predicts that chemicals that bind to these receptors should be further tested for adverse effects. Contrary to BPA, 17β-estradiol, the natural ligand of estrogen receptors, did not induce apoptosis; if anything, 17β-estradiol diminished it [33]. These data show that different estrogen receptor ligands may have opposite effects on β-cells, suggesting that ligand binding to estrogen receptors does not predict its effect on apoptosis, at least in β-cells. In line with previous in vivo [48,49] and in vitro [21,25,33] studies, we observed that BPA induced ROS production, which suggests that ROS generation might be considered an important key event that could predict β-cell death.

Our experiments indicate that TBT also induces apoptosis. In animal models, Zuo and colleagues showed that oral administration of 0.5, 5, and 50 μg/kg TBT every three days for 45 days induced apoptosis while inhibiting proliferation of mouse islet cells [50]. In rats, TBT effects on cell viability varied depending on the model studied. A TBT dose as low as 10 nM decreased cell viability by 20% in isolated rat islets [27], whereas doses up to 200 nM did not affect viability in the rat insulinoma cell line RINm5F [22,29]. Yet, exposure to 500 nM TBT for 24 h markedly increased the number of apoptotic cells in the RINm5F cell line [22]. In line with the results observed in mouse [50] and rat islets [27], here we show that doses from 1 to 200 nM reduced cell viability by inducing apoptosis in two cell lines, EndoC-βH1 and INS-1E, and in dispersed mouse islets. TBT is an agonist of PPARγ and RXR [34,35] and our results indicate that the apoptotic effect of TBT in β-cells is mediated by these receptors because it is abolished by a PPARγ antagonist. Like BPA, TBT also increased ROS levels, supporting a role for ROS production as a key event in the pathway to MDC-induced β-cell apoptosis.

Besides BPA and TBT, exposure to the other four MDCs did not change β-cell viability at the doses tested. One of these EDCs, PFOA, did not change ROS production, confirming that assessment of ROS levels is a good predictor of apoptosis in our model. Regarding cell viability, our findings are in line with the studies available for PFOA and TCS. Following 48 h exposure, 1 μM PFOA did not induce apoptosis in rat RIN-m5F β-cells; only doses higher than 100 μM (up to 500 μM) decreased cell viability due to apoptosis [51]. Of note, we also found that higher concentrations of PFOA (20 to 200 μM) induced apoptosis in EndoC-βH1 and INS-1E cells (data not shown). Regarding TCS, Ajao et al. found that 17 and 35 μM TCS induced β-cell death by necrosis in the mouse β-cell line MIN-6, whereas 7 μM TCS did not change MIN-6 viability [52]. This agrees with our results, in which 1 μM TCS did not change β-cell viability. Despite the lack of evidence in pancreatic β-cells, it has been shown that exposure to high doses of TPP and DDE (>10 μM) induced apoptosis in other cells and tissues: TPP in hepatocytes, epithelial tumoral JEG-3 cells from placenta, and murine BV-2 microglia [53–55]; and DDE in Sertoli cells [56,57]. Exposure to high concentrations of *p,p’*-DDT (>150 μM) for 24 h decreased viability in NES2Y human pancreatic β-cell line [58]. Lack of publications with these EDCs at low concentrations does not allow comparison with the present findings.

Finally, our present results indicate that EndoC-βH1 cells are a robust human β-cell model for identifying MDCs that alter β-cell viability. Furthermore, the methods used herein to measure cell viability and/or apoptosis are easy to use, particularly the MTT assay, and appear to be suitable for testing cell death in response to MDCs. Assessment of ROS production as a key event correctly predicted β-cell death, suggesting that it could be a useful surrogate for measurement of cell viability.

### 3.2. β-cell function tests

INS-1E have been used for studies of β-cell secretion for the last two decades. They present a similar GSIS to rat islets with a maximal increase of about 6-fold at 15 mM glucose and a 50% effective concentration about 10 mM [12]. Its electrical activity is, however, different to that described for islet cells and it is saturated at a concentration of 7.5 mM glucose [12]. EndoC-βH1 is an appropriate model to study GSIS; these cells release more insulin at basal and stimulatory (20 mM) glucose levels than INS-1 832/13 cells and their response to high glucose is comparable to human islets, *i.e*., a 2- to 2.5-fold increase in insulin secretion upon stimulation [14]. We also report a 2- to 3-fold increase upon stimulation with 20 mM glucose, which is similar to what has been published in the literature [13,14,41,42], indicating that our cells respond adequately to glucose. Conversely, due to technical issues, we were unable to use INS-1E cells for analyses of GSIS and insulin content. Similarly, an earlier study demonstrated that the rat β-cell line INS-1 832/13 seemed to lack certain characteristics that made it unsuitable as a screening system for diabetogenic contaminants [26].

Our results indicate that PFOA was the MDC that produced the most changes in insulin secretion in EndoC-βH1 cells at basal and glucose-stimulating levels. This MDC induced a decrease in GSIS at very low doses, within the pM and nM range while no effect was observed at 1 μM. Interestingly, neither apoptosis nor ROS production were induced upon exposure to 1 nM and 1 μM PFOA. These data suggest that the decrease in insulin release shown here is not due to a cytotoxic action of PFOA. We did not test higher PFOA concentrations because they are not within the range of human exposure. Nonetheless, a previous study demonstrated that higher concentrations of PFOA, above 10 μM, increased insulin release in the mouse-derived β-TC-6 β-cell line through the G-protein coupled receptor GPR40 [59]. Studies in animal models indicate that PFOA alters glucose homeostasis. In adult male mice, oral administration of PFOA 1.25 mg/kg for 28 days increased fasting blood glucose levels and decreased hepatic glycogen and glucose content, but did not influence insulin blood levels [60]. Higher PFOA doses (5 mg/kg) induced hyperinsulinemia in the fasted state [61]. In CD-1 mice offspring exposed in utero, low concentrations of PFOA (0.01-0.1mg/kg) increased weight gain as well as insulin and leptin levels in blood [62]. These alterations in insulin blood levels could be due to an indirect rather than a direct effect on β-cells. Unfortunately, we could not find any evidence of direct effects of low doses of PFOA on insulin secretion so that we could compare with our data. Although our results need further confirmation, they point to PFOA as a potential diabetogenic MDC.

As a key event with the potential to predict alterations in β-cell function, we measured expression of five key genes involved in β-cell identity (*PDX1* and *MAFA*) and function (*INS, GLUT2* and *GCK*). None of those genes changed with PFOA treatment, indicating that these genes may not be good candidates to predict alterations in β-cell function.

The other MDCs that changed GSIS were TBT and TPP at the highest dose tested and DDE at 10 nM and 1 μM. Previous work suggested that TBT at 100 and 200 nM increased GSIS in RIN-m5F cells and in human islets [29], which is in the same concentration range shown in our work. This increase in GSIS promoted by 200 nM TBT correlates with an increase in the expression of *GLUT2* (a gene that encodes the glucose transporter 2), which could increase glucose uptake. Notably, TBT decreased *MAFA* expression even at low concentrations in EndoC-βH1 and at 200 nM in INS-1E. MAFA controls glucose-responsive transcription of insulin and other genes in β-cells [63]. Insulin content decreased with 200 nM TBT, which might be, in part, due to *MAFA* downregulation. Insulin content, however, was not diminished upon treatment with TBT doses that decreased *MAFA* expression. On the contrary, 20 nM TBT increased insulin content. Then, even though our findings regarding *MAFA* expression are of interest, further research is required to link its downregulation to changes in insulin content.

Little is known about TPP effects on insulin and glucose homeostasis. In other metabolism-related cell systems, such as differentiated 3T3-L1 adipocytes, TPP exposure increased glucose uptake and lipolysis [64]. In animal models, perinatal exposure to TPP accelerated type 2 diabetes onset [65] and a mix of organophosphate flame retardants, including TPP, altered glucose homeostasis in an ERα-dependent manner [66]. Mice exposed to TPP during fetal development presented impaired glucose tolerance and increased insulin levels [67]. Here we show that exposure to 1 μM TPP induced an increase in insulin release, despite the lack of changes on gene expression at 1 nM or 1 μM. As this is the first time that a direct effect of TPP on β-cells is shown, we take our findings carefully until they can be further validated.

DDE is a metabolite of the persistent endocrine disruptor DDT. Previous experiments performed in INS-1E cells showed that exposure to non-lethal concentrations of DDT or DDE changed insulin secretion and content/expression [58,68]. Upon 48 h treatment, low concentrations of DDT (1 fM to 1 μM) decreased GSIS in a non-monotonic manner, while all doses tested diminished intracellular insulin content [68]. In addition, INS-1E cells exposed for 1 month to a non-lethal concentration (10 μM) of DDT or DDE had lower mRNA levels of *Ins1* and *Ins2* as well as reduced expression of proinsulin and insulin [58]. On the other hand, DDE exposure increased insulin secretion in β-TC-6 cells [69], which agrees with our results that suggest an increase of GSIS at 10 nM and 1 μM DDE (insulin content remained unchanged in all concentrations tested).

The findings described in the present work highly suggest that the EndoC-βH1 cell line is a valid human β-cell model to study the effect of MDCs on cell viability and function. We also present the rat INS-1E cells may represent a valuable, alternative model for testing the effect of MDCs on cell viability because results mirror that obtained with EndoC-βH1 cells. Altogether, we conclude that EndoC-βH1 may represent a valuable model of human β-cells for screening and testing procedures to identify MDC with diabetogenic activity. MTT assay and staining with DNA-binding dyes Hoechst 33342 and propidium iodide represent useful test methods to evaluate cell viability, while assessment of ROS production is a potential substitute for cell viability assays. Finally, measurement of GSIS seems a more reliable test method to investigate MDC effects on β-cell function.

## 4. Materials and Methods

### 4.1. Chemicals

Chemicals used herein were purchased as follows: BPA (Cat. No. 239658), TBT (Cat. No. T50202), PFOA (Cat. No. 77262), TPP (Cat. No. 241288), TCS (Cat. No. PHR1338), and DDE (Cat. No. 123897) were obtained from Sigma-Aldrich (Barcelona, Spain). All stock solutions were weekly prepared by dissolution in 100% cell-culture grade, sterile-filtered DMSO (Sigma-Aldrich; Cat No D2650) and stored at −20°C between uses. ICI 182,780 (Cat. No. 1047) and T0070907 (Cat. No. HY-13202) were obtained from Tocris Cookson Ltd (Avonmouth, UK) and MedChem Express (New Jersey, USA), respectively. Due to its short half-life, T0070907 was redosed every 8-10 h.

### 4.2. Culture of INS-1E and EndoC-βH1 cells

Rat insulin-producing INS-1E cells (RRID: CVCL_0351, kindly provided by Dr. C. Wollheim, Department of Cell Physiology and Metabolism, University of Geneva, Geneva, Switzerland) were cultured in RPMI 1640 GlutaMAX-I, 10 mM HEPES, 1 mM sodium pyruvate, 5% fetal bovine serum, 50 μM 2-mercaptoethanol, 50 units/ml penicillin, and 50 mg/ml streptomycin [70]. Human insulin-producing EndoC-βH1 cells (RRID: CVCL_L909, Univercell-Biosolutions, France), which grow attached to Matrigel/fibronectin-coated plates, were cultured in DMEM containing 5.6 mM glucose, 10 mM nicotinamide, 6.7 ng/ml selenite, 5.5 μg/ml transferrin, 50 μM 2-mercaptoethanol, 2% BSA fatty acid free, 100 U/ml penicillin, and 100 μg/ml streptomycin [13]. For treatments in EndoC-βH1 cells, 2% FBS is added to the culture medium. Both cell lines were kept at 37°C in a humidified atmosphere of 95% O_2_ and 5% CO_2_.

### 4.3. Isolation and dispersion of mouse islets

Adult male mice (12-14-weeks-old), which were kept under standard housing conditions (12 h light/dark cycle, food *ad libitum*), were sacrificed and pancreatic islets were isolated as described [71]. Islets were dispersed into single cells and cultured in polylysine-coated plates as described before [72,73]. Dispersed cells were kept at 37 °C in a humidified atmosphere of 95% O_2_ and 5% CO_2_ and used within 48 h of culture. Experimental procedures were performed according to the Spanish Royal Decree 1201/2005 and the European Community Council directive 2010/63/EU. The ethical committee of Miguel Hernandez University reviewed and approved the methods used herein (approvals ID: UMH-IB-AN-01-14 and UMH-IB-AN-02-14).

### 4.4. Assessment of cell viability by MTT assay

The MTT compound (3-(4,5-dimethylthiazol-2-yl)-2,5-diphenyltetrazolium bromide) (Sigma-Aldrich) was dissolved in RPMI 1640 without phenol red. MTT was added to each well (0.5 mg/ml) and incubated for 3 h at 37°C. After incubation, the supernatant was removed by aspiration and 100 μl DMSO was added to dissolve formazan crystals. The absorbance was measured at 595 nm using an iMark™ Microplate Absorbance Reader (Bio-Rad, Hercules, CA) and the percentage of cell viability was calculated.

### 4.5. Assessment of cell viability by DNA-binding dyes

Percentage of living, apoptotic and necrotic cells was determined after staining with DNA-binding dyes Hoechst 33342 (HO) and propidium iodide (PI) as described [70,74]. Two different observers, one of them being unaware of sample identity to avoid bias, counted a minimum of 500 cells per experimental condition (agreement between results from both observers was >90%). Results are expressed as percentage of apoptosis.

### 4.6. Caspase 3/7 activity assay

Caspase 3/7 activity was determined using the Caspase-Glo^®^ 3/7 assay (Promega, Madison, WI, USA) according to the manufacturer’s instructions. Briefly, upon 48 h treatment in 100 μl culture medium, cells were incubated with 100 μl Caspase-Glo® 3/7 reagent at room temperature for 1 h before recording luminescence with a POLASTAR plate reader (BMG Labtech, Germany).

### 4.7. DCF assay

ROS production was measured using the fluorescent probe 2’,7’-dichlorofluorescein di-acetate (DCF; Sigma-Aldrich). Cells were seeded in 96-well black plates and, upon treatment, were loaded with 10□μM DCF for 30□min at 37°C. DCF fluorescence was quantified in a POLASTAR plate reader (BMG Labtech, Germany). Data are expressed as DCF fluorescence corrected by total protein. Menadione (15 μM for 90 minutes) was used as a positive control.

### 4.8. Glucose-stimulated insulin secretion

Cells were preincubated in Krebs-Ringer buffer (115 mM NaCl, 5 mM KCl, 1 mM MgCl_2_, 1 mM CaCl_2_, 24 mM NaHCO_3_, 10 mM HEPES pH 7.4, and 0.1% BSA) for 1 h before glucose stimulation. At the end of this incubation, cells were sequentially stimulated with low (0 mM) and then high glucose (20 mM) for 40 min (each stimulation). After each stimulatory period, the incubation medium was collected, placed onto ice, and centrifuged at 700 *g*, 5 min at 4 °C. The supernatant was transferred into a fresh tube and stored at −20 °C until insulin measurements. For insulin content, cells were lysed in 100 μl of cell lysis solution (137 mM NaCl, 1% Glycerol, 0.1% Triton X100, 2 mM EGTA, 20 mM Tris pH 8.0, and protease inhibitor cocktail). Cell lysates were centrifuged at 700 *g*, 5 min at 4 °C. The supernatant was transferred into a fresh tube and stored at −20 °C until insulin measurements. Insulin release and content were measured using a human insulin ELISA kit (Mercodia, Uppsala, Sweden).

The amount of secreted insulin secretion as % of total insulin was calculated as previously described [42]. Data are normalized to insulin secretion at high glucose in vehicle-treated cells.

### 4.9. RNA extraction and real-time PCR

Total RNA was isolated using the RNeasy Micro Kit (Qiagen) and poly(A)^+^ mRNA extraction was performed using Dynabeads mRNA DIRECT kit (Invitrogen) in accordance with the manufacturer’s instructions. cDNA synthesis was performed using the High-Capacity cDNA Reverse Transcription Kit (Applied Biosystems). Quantitative PCR was carried out using the CFX96 Real Time System (Bio-Rad) as described [75]. *Gapdh* and β-actin were used as housekeeping genes for rat and human samples, respectively. The CFX Manager Version 1.6 (Bio-Rad) was used to analyze the values, which were expressed as relative expression. The primers used herein are listed in Supplemental Table 1.

### 4.10. Data analysis

The GraphPad Prism 7.0 software (GraphPad Software, La Jolla, CA, USA) was used for statistical analyses. Data are presented as the mean ± SEM. Statistical analyses were performed using Student’s t-test, one-way ANOVA, or two-way ANOVA. *p* values ≤ 0.05 were considered statistically significant.

## Acknowledgements

The authors thank Beatriz Bonmati Botella, Maria Luisa Navarro, and Salomé Ramon for their excellent technical assistance. Graphical Abstract was adapted from “Mechanisms behind the induction of trained Immunity”, by BioRender.com (2022). Retrieved from https://app.biorender.com/biorender-templates.

## Author contributions

R.S.d.S.: Conceptualization, Supervision, Visualization, Investigation, Formal analysis, Writing – original draft & editing; R.M.M-G.: Investigation, Formal analysis, Writing – review & editing; I.B-C.: Investigation, Formal analysis, Writing – review & editing; L.M.: Investigation, Formal analysis, Writing – review & editing; A.N.: Conceptualization, Supervision, Visualization, Resources, Funding acquisition, Writing – original draft & editing, Project administration. All authors have read and given approval to the final version of the manuscript.

## Institutional Review Board Statement

The study was conducted according to the guidelines of the Declaration of Helsinki and approved by the Spanish Royal Decree 1201/2005 and the European Community Council directive 2010/63/EU. The ethical committee of Miguel Hernandez University reviewed and approved the methods used herein (approvals ID: UMH-IB-AN-01-14 and UMH-IB-AN-02-14).

## References

1. Sun, H.; Saeedi, P.; Karuranga, S.; Pinkepank, M.; Ogurtsova, K.; Duncan, B.B.; Stein, C.; Basit, A.; Chan, J.C.N.; Mbanya, J.C.; et al. IDF Diabetes Atlas: Global, regional and country-level diabetes prevalence estimates for 2021 and projections for 2045. Diabetes Res. Clin. Pract. 2022, 183, 109119, doi:10.1016/j.diabres.2021.109119.

2. Eizirik, D.L.; Pasquali, L.; Cnop, M. Pancreatic β-cells in type 1 and type 2 diabetes mellitus: different pathways to failure. Nat. Rev. Endocrinol. 2020, 16, 349–362, doi:10.1038/s41574-020-0355-7.

3. Marroqui, L.; Perez-Serna, A.A.; Babiloni-Chust, I.; Dos Santos, R.S. Type I interferons as key players in pancreatic β-cell dysfunction in type 1 diabetes. Int. Rev. Cell Mol. Biol. 2021, 359, 1–80, doi:10.1016/bs.ircmb.2021.02.011.

4. Howard, S.G. Developmental Exposure to Endocrine Disrupting Chemicals and Type 1 Diabetes Mellitus. Front. Endocrinol. (Lausanne). 2018, 9, doi:10.3389/fendo.2018.00513.

5. Esser, N.; Utzschneider, K.M.; Kahn, S.E. Early beta cell dysfunction vs insulin hypersecretion as the primary event in the pathogenesis of dysglycaemia. Diabetologia 2020, 63, 2007–2021, doi:10.1007/s00125-020-05245-x.

6. Corkey, B.E. Banting lecture 2011: Hyperinsulinemia: Cause or consequence? Diabetes 2012, 61.

7. Alonso-Magdalena, P.; Quesada, I.; Nadal, A. Endocrine disruptors in the etiology of type 2 diabetes mellitus. Nat. Rev. Endocrinol. 2011, 7, 346–353, doi:10.1038/nrendo.2011.56.

8. Heindel, J.J.; Blumberg, B.; Cave, M.; Machtinger, R.; Mantovani, A.; Mendez, M.A.; Nadal, A.; Palanza, P.; Panzica, G.; Sargis, R.; et al. Metabolism disrupting chemicals and metabolic disorders. Reprod. Toxicol. 2017, 68, doi:10.1016/j.reprotox.2016.10.001.

9. Mimoto, M.S.; Nadal, A.; Sargis, R.M. Polluted Pathways: Mechanisms of Metabolic Disruption by Endocrine Disrupting Chemicals. Curr. Environ. Heal. reports 2017, 4, doi:10.1007/s40572-017-0137-0.

10. Nadal, A.; Quesada, I.; Tudurí, E.; Nogueiras, R.; Alonso-Magdalena, P. Endocrine-disrupting chemicals and the regulation of energy balance. Nat. Rev. Endocrinol. 2017, 13, doi:10.1038/nrendo.2017.51.

11. Legler, J.; Zalko, D.; Jourdan, F.; Jacobs, M.; Fromenty, B.; Balaguer, P.; Bourguet, W.; Kos, V.M.; Nadal, A.; Beausoleil, C.; et al. The GOLIATH project: Towards an internationally harmonised approach for testing metabolism disrupting compounds. Int. J. Mol. Sci. 2020, 21, 3480, doi:10.3390/ijms21103480.

12. Merglen, A.; Theander, S.; Rubi, B.; Chaffard, G.; Wollheim, C.B.; Maechler, P. Glucose Sensitivity and Metabolism-Secretion Coupling Studied during Two-Year Continuous Culture in INS-1E Insulinoma Cells. Endocrinology 2004, 145, 667–678, doi:10.1210/en.2003-1099.

13. Ravassard, P.; Hazhouz, Y.; Pechberty, S.; Bricout-Neveu, E.; Armanet, M.; Czernichow, P.; Scharfmann, R. A genetically engineered human pancreatic β cell line exhibiting glucose-inducible insulin secretion. J. Clin. Invest. 2011, 121, 3589–97, doi:10.1172/JCI58447.

14. Andersson, L.E.; Valtat, B.; Bagge, A.; Sharoyko, V. V.; Nicholls, D.G.; Ravassard, P.; Scharfmann, R.; Spégel, P.; Mulder, H. Characterization of Stimulus-Secretion Coupling in the Human Pancreatic EndoC-βH1 Beta Cell Line. PLoS One 2015, 10, e0120879, doi:10.1371/journal.pone.0120879.

15. Vandenberg, L.N.; Chahoud, I.; Heindel, J.J.; Padmanabhan, V.; Paumgartten, F.J.R.; Schoenfelder, G. Urinary, circulating, and tissue biomonitoring studies indicate widespread exposure to bisphenol A. Environ. Health Perspect. 2010, 118, 1055–1070, doi:10.1289/ehp.0901716.

16. Kannan, K.; Senthilkumar, K.; Giesy, J.P. Occurrence of butyltin compounds in human blood. Environ. Sci. Technol. 1999, 33, doi:10.1021/es990011w.

17. Göckener, B.; Weber, T.; Rüdel, H.; Bücking, M.; Kolossa-Gehring, M. Human biomonitoring of per-and polyfluoroalkyl substances in German blood plasma samples from 1982 to 2019. Environ. Int. 2020, 145, doi:10.1016/j.envint.2020.106123.

18. Dodson, R.E.; Eede, N. Van Den; Covaci, A.; Perovich, L.J.; Brody, J.G.; Rudel, R.A. Urinary biomonitoring of phosphate flame retardants: Levels in california adults and recommendations for future studies. Environ. Sci. Technol. 2014, 48, doi:10.1021/es503445c.

19. Tschersich, C.; Murawski, A.; Schwedler, G.; Rucic, E.; Moos, R.K.; Kasper-Sonnenberg, M.; Koch, H.M.; Brüning, T.; Kolossa-Gehring, M. Bisphenol A and six other environmental phenols in urine of children and adolescents in Germany – human biomonitoring results of the German Environmental Survey 2014–2017 (GerES V). Sci. Total Environ. 2021, 763, doi:10.1016/j.scitotenv.2020.144615.

20. Henríquez-Hernández, L.A.; Ortiz-Andrelluchi, A.; Álvarez-Pérez, J.; Acosta-Dacal, A.; Zumbado, M.; Martínez-González, M.A.; Boada, L.D.; Salas-Salvadó, J.; Luzardo, O.P.; Serra-Majem, L. Human biomonitoring of persistent organic pollutants in elderly people from the Canary Islands (Spain): A temporal trend analysis from the PREDIMED and PREDIMED-Plus cohorts. Sci. Total Environ.2021, 758, doi:10.1016/j.scitotenv.2020.143637.

21. Carchia, E.; Porreca, I.; Almeida, P.J.; D’Angelo, F.; Cuomo, D.; Ceccarelli, M.; De Felice, M.; Mallardo, M.; Ambrosino, C. Evaluation of low doses BPA-induced perturbation of glycemia by toxicogenomics points to a primary role of pancreatic islets and to the mechanism of toxicity. Cell Death Dis. 2015, 6, e1959–e1959, doi:10.1038/cddis.2015.319.

22. Huang, C.F.; Yang, C.Y.; Tsai, J.R.; Wu, C.T.; Liu, S.H.; Lan, K.C. Low-dose tributyltin exposure induces an oxidative stress-triggered JNK-related pancreatic β-cell apoptosis and a reversible hypoinsulinemic hyperglycemia in mice. Sci. Rep. 2018, 8, doi:10.1038/s41598-018-24076-w.

23. Alonso-Magdalena, P.; Ropero, A.B.; Carrera, M.P.; Cederroth, C.R.; Baquié, M.; Gauthier, B.R.; Nef, S.; Stefani, E.; Nadal, A. Pancreatic insulin content regulation by the Estrogen receptor ERα. PLoS One 2008, 3, doi:10.1371/journal.pone.0002069.

24. Marroqui, L.; Martinez-Pinna, J.; Castellano-Muñoz, M.; dos Santos, R.S.; Medina-Gali, R. M.; Soriano, S.; Quesada, I.; Gustafsson, J.A.; Encinar, J.A.; Nadal, A. Bisphenol-S and Bisphenol-F alter mouse pancreatic β-cell ion channel expression and activity and insulin release through an estrogen receptor ERβ mediated pathway. Chemosphere 2021, 265, doi:10.1016/j.chemosphere.2020.129051.

25. Lin, Y.; Sun, X.; Qiu, L.; Wei, J.; Huang, Q.; Fang, C.; Ye, T.; Kang, M.; Shen, H.; Dong, S. Exposure to bisphenol A induces dysfunction of insulin secretion and apoptosis through the damage of mitochondria in rat insulinoma (INS-1) cells. Cell Death Dis. 2013, 4, e460–e460, doi:10.1038/cddis.2012.206.

26. Hectors, T.L.M.; Vanparys, C.; Pereira-Fernandes, A.; Martens, G.A.; Blust, R. Evaluation of the INS-1 832/13 Cell Line as a Beta-Cell Based Screening System to Assess Pollutant Effects on Beta-Cell Function. PLoS One 2013, 8, e60030, doi:10.1371/journal.pone.0060030.

27. Ghaemmaleki, F.; Mohammadi, P.; Baeeri, M.; Navaei-Nigjeh, M.; Abdollahi, M.; Mostafalou, S. Estrogens counteract tributyltin-induced toxicity in the rat islets of Langerhans. Heliyon 2020, 6, e03562, doi:10.1016/j.heliyon.2020.e03562.

28. Soriano, S.; Alonso-Magdalena, P.; García-Arévalo, M.; Novials, A.; Muhammed, S.J.; Salehi, A.; Gustafsson, J. A.; Quesada, I.; Nadal, A. Rapid insulinotropic action of low doses of Bisphenol-A on mouse and human islets of Langerhans: Role of estrogen receptor β. PLoS One 2012, 7, e31109, doi:10.1371/journal.pone.0031109.

29. Chen, Y.W.; Lan, K.C.; Tsai, J.R.; Weng, T.I.; Yang, C.Y.; Liu, S.H. Tributyltin exposure at noncytotoxic doses dysregulates pancreatic β-cell function in vitro and in vivo. Arch. Toxicol. 2017, 91, 3135–3144, doi:10.1007/s00204-017-1940-y.

30. Denizot, F.; Lang, R. Rapid colorimetric assay for cell growth and survival. J. Immunol. Methods 1986, 89, 271–277, doi:10.1016/0022-1759(86)90368-6.

31. Mosmann, T. Rapid colorimetric assay for cellular growth and survival: Application to proliferation and cytotoxicity assays. J. Immunol. Methods 1983, 65, doi: 10.1016/0022-1759(83)90303-4.

32. Hoorens, A.; Van De Casteele, M.; Klöppel, G.; Pipeleers, D. Glucose promotes survival of rat pancreatic β cells by activating synthesis of proteins which suppress a constitutive apoptotic program. J. Clin. Invest. 1996, 98, 1568–1574, doi:10.1172/JCI118950.

33. Babiloni-Chust, I.; dos Santos, R.S.; Medina-Gali, R.M.; Perez-Serna, A.A.; Encinar, J.-A.; Martinez-Pinna, J.; Gustafsson, J.-A.; Marroqui, L.; Nadal, A. G protein-coupled estrogen receptor activation by Bisphenol-A disrupts protection from apoptosis conferred by estrogen receptors ERα and ERβ in pancreatic beta cells. bioRxiv 2022, 2022.01.31.478472, doi:10.1101/2022.01.31.478472.

34. Gru□n, F.; Blumberg, B. Environmental Obesogens: Organotins and Endocrine Disruption via Nuclear Receptor Signaling. Endocrinology 2006, 147, s50–s55, doi:10.1210/en.2005-1129.

35. le Maire, A.; Grimaldi, M.; Roecklin, D.; Dagnino, S.; Vivat Hannah, V.; Balaguer, P.; Bourguet, W. Activation of RXR–PPAR heterodimers by organotin environmental endocrine disruptors. EMBO Rep. 2009, 10, 367–373, doi:10.1038/embor.2009.8.

36. Scharfmann, R.; Staels, W.; Albagli, O. The supply chain of human pancreatic β cell lines. J. Clin. Invest. 2019, 129, 3511–3520, doi:10.1172/JCI129484.

37. Scharfmann, R.; Didiesheim, M.; Richards, P.; Chandra, V.; Oshima, M.; Albagli, O. Mass production of functional human pancreatic β-cells: why and how? Diabetes, Obes. Metab. 2016, 18, 128–136, doi:10.1111/dom.12728.

38. Lawlor, N.; Márquez, E.J.; Orchard, P.; Narisu, N.; Shamim, M.S.; Thibodeau, A.; Varshney, A.; Kursawe, R.; Erdos, M.R.; Kanke, M.; et al. Multiomic Profiling Identifies cis-Regulatory Networks Underlying Human Pancreatic β Cell Identity and Function. Cell Rep. 2019, 26, 788–801.e6, doi:10.1016/j.celrep.2018.12.083.

39. Ryaboshapkina, M.; Saitoski, K.; Hamza, G.M.; Jarnuczak, A.F.; Berthault, C.; Sengupta, K.; Underwood, C.R.; Andersson, S.; Scharfmann, R. Characterization of the secretome, transcriptome and proteome of human β cell line EndoC-βH1. bioRxiv 2021, 2021.09.09.459582, doi:10.1101/2021.09.09.459582.

40. Gurgul-Convey, E.; Kaminski, M.T.; Lenzen, S. Physiological characterization of the human EndoC-βH1 β-cell line. Biochem. Biophys. Res. Commun. 2015, 464, 13–19, doi:10.1016/j.bbrc.2015.05.072.

41. Hastoy, B.; Godazgar, M.; Clark, A.; Nylander, V.; Spiliotis, I.; van de Bunt, M.; Chibalina, M. V; Barrett, A.; Burrows, C.; Tarasov, A.I.; et al. Electrophysiological properties of human beta-cell lines EndoC-βH1 and −βH2 conform with human beta-cells. Sci. Rep. 2018, 8, 16994, doi:10.1038/s41598-018-34743-7.

42. Tsonkova, V.G.; Sand, F.W.; Wolf, X.A.; Grunnet, L.G.; Ringgaard, A.K.; Ingvorsen, C.; Winkel, L.; Kalisz, M.; Dalgaard, K.; Bruun, C.; et al. The EndoC-βH1 cell line is a valid model of human beta cells and applicable for screenings to identify novel drug target candidates. Mol. Metab. 2018, 8, 144–157, doi:10.1016/j.molmet.2017.12.007.

43. Diedisheim, M.; Oshima, M.; Albagli, O.; Huldt, C.W.; Ahlstedt, I.; Clausen, M.; Menon, S.; Aivazidis, A.; Andreasson, A.C.; Haynes, W.G.; et al. Modeling human pancreatic beta cell dedifferentiation. Mol. Metab. 2018, 10, 74–86, doi:10.1016/j.molmet.2018.02.002.

44. Colli, M.L.; Ramos-Rodríguez, M.; Nakayasu, E.S.; Alvelos, M.I.; Lopes, M.; Hill, J.L.E.; Turatsinze, J.V.; Coomans de Brachène, A.; Russell, M.A.; Raurell-Vila, H.; et al. An integrated multi-omics approach identifies the landscape of interferon-α-mediated responses of human pancreatic beta cells. Nat. Commun. 2020, 11, 2584, doi:10.1038/s41467-020-16327-0.

45. Marroqui, L.; Santin, I.; Dos Santos, R.S.; Marselli, L.; Marchetti, P.; Eizirik, D.L. BACH2, a Candidate Risk Gene for Type 1 Diabetes, Regulates Apoptosis in Pancreatic β-Cells via JNK1 Modulation and Crosstalk With the Candidate Gene PTPN2. Diabetes 2014, 63, 2516–2527, doi:10.2337/db13-1443.

46. Dos Santos, R.S.; Marroqui, L.; Grieco, F.A.; Marselli, L.; Suleiman, M.; Henz, S.R.; Marchetti, P.; Wernersson, R.; Eizirik, D.L. Protective role of complement C3 against cytokine-mediated β-cell apoptosis. Endocrinology 2017, 158, 2503–2521, doi:10.1210/en.2017-00104.

47. Le Magueresse-Battistoni, B.; Multigner, L.; Beausoleil, C.; Rousselle, C. Effects of bisphenol A on metabolism and evidences of a mode of action mediated through endocrine disruption. Mol. Cell. Endocrinol. 2018, 475, 74–91, doi:10.1016/j.mce.2018.02.009.

48. Wei, J.; Lin, Y.; Li, Y.; Ying, C.; Chen, J.; Song, L.; Zhou, Z.; Lv, Z.; Xia, W.; Chen, X.; et al. Perinatal Exposure to Bisphenol A at Reference Dose Predisposes Offspring to Metabolic Syndrome in Adult Rats on a High-Fat Diet. Endocrinology 2011, 152, 3049–3061, doi:10.1210/en.2011-0045.

49. Bansal, A.; Rashid, C.; Xin, F.; Li, C.; Polyak, E.; Duemler, A.; van der Meer, T.; Stefaniak, M.; Wajid, S.; Doliba, N.; et al. Sex-and dose-specific effects of maternal bisphenol A exposure on pancreatic islets of first-and second-generation adult mice offspring. Environ. Health Perspect. 2017, 125, doi:10.1289/EHP1674.

50. Zuo, Z.; Wu, T.; Lin, M.; Zhang, S.; Yan, F.; Yang, Z.; Wang, Y.; Wang, C. Chronic exposure to tributyltin chloride induces pancreatic islet cell apoptosis and disrupts glucose homeostasis in male mice. Environ. Sci. Technol. 2014, 48, 5179–5186, doi:10.1021/es404729p.

51. Suh, K.S.; Choi, E.M.; Kim, Y.J.; Hong, S.M.; Park, S.Y.; Rhee, S.Y.; Oh, S.; Kim, S.W.; Pak, Y.K.; Choe, W.; et al. Perfluorooctanoic acid induces oxidative damage and mitochondrial dysfunction in pancreatic β-cells. Mol. Med. Rep. 2017, 15, 3871–3878, doi:10.3892/mmr.2017.6452.

52. Ajao, C.; Andersson, M.A.; Teplova, V. V.; Nagy, S.; Gahmberg, C.G.; Andersson, L.C.; Hautaniemi, M.; Kakasi, B.; Roivainen, M.; Salkinoja-Salonen, M. Mitochondrial toxicity of triclosan on mammalian cells. Toxicol. Reports2015, 2, 624–637, doi:10.1016/j.toxrep.2015.03.012.

53. Bowen, C.; Childers, G.; Perry, C.; Martin, N.; McPherson, C.A.; Lauten, T.; Santos, J.; Harry, G.J. Mitochondrial-related effects of pentabromophenol, tetrabromobisphenol A, and triphenyl phosphate on murine BV-2 microglia cells. Chemosphere 2020, 255, 126919, doi:10.1016/j.chemosphere.2020.126919.

54. Wang, Y.; Hong, J.; Shi, M.; Guo, L.; Liu, L.; Tang, H.; Liu, X. Triphenyl phosphate disturbs the lipidome and induces endoplasmic reticulum stress and apoptosis in JEG-3 cells. Chemosphere 2021, 275, 129978, doi:10.1016/j.chemosphere.2021.129978.

55. Wang, X.; Li, F.; Liu, J.; Li, Q.; Ji, C.; Wu, H. New insights into the mechanism of hepatocyte apoptosis induced by typical organophosphate ester: An integrated in vitro and in silico approach. Ecotoxicol. Environ. Saf. 2021, 219, 112342, doi:10.1016/j.ecoenv.2021.112342.

56. Song, Y.; Liang, X.; Hu, Y.; Wang, Y.; Yu, H.; Yang, K. p,p’-DDE induces mitochondria-mediated apoptosis of cultured rat Sertoli cells. Toxicology 2008, 253, 53–61, doi:10.1016/j.tox.2008.08.013.

57. Shi, Y.; Song, Y.; Wang, Y.; Liang, X.; Hu, Y.; Guan, X.; Cheng, J.; Yang, K. p,p’-DDE Induces Apoptosis of Rat Sertoli Cells via a FasL-Dependent Pathway. J. Biomed. Biotechnol. 2009, 2009, 1–11, doi:10.1155/2009/181282.

58. Pavlíková, N.; Daniel, P.; Šrámek, J.; Jelínek, M.; Šrámková, V.; Němcová, V.; Balušíková, K.; Halada, P.; Kovář, J. Upregulation of vitamin D-binding protein is associated with changes in insulin production in pancreatic beta-cells exposed to p,p’-DDT and p,p’-DDE. Sci. Rep. 2019, 9, 18026, doi:10.1038/s41598-019-54579-z.

59. Qin, W.P.; Cao, L.Y.; Li, C.H.; Guo, L.H.; Colbourne, J.; Ren, X.M. Perfluoroalkyl Substances Stimulate Insulin Secretion by Islet β Cells via G Protein-Coupled Receptor 40. Environ. Sci. Technol. 2020, 54, 3428–3436, doi:10.1021/acs.est.9b07295.

60. Zheng, F.; Sheng, N.; Zhang, H.; Yan, S.; Zhang, J.; Wang, J. Perfluorooctanoic acid exposure disturbs glucose metabolism in mouse liver. Toxicol. Appl. Pharmacol. 2017, 335, doi:10.1016/j.taap.2017.09.019.

61. Wu, X.; Xie, G.; Xu, X.; Wu, W.; Yang, B. Adverse bioeffect of perfluorooctanoic acid on liver metabolic function in mice. Environ. Sci. Pollut. Res. 2018, 25, 4787–4793, doi:10.1007/s11356-017-0872-7.

62. Hines, E.P.; White, S.S.; Stanko, J.P.; Gibbs-Flournoy, E.A.; Lau, C.; Fenton, S.E. Phenotypic dichotomy following developmental exposure to perfluorooctanoic acid (PFOA) in female CD-1 mice: Low doses induce elevated serum leptin and insulin, and overweight in mid-life. Mol. Cell. Endocrinol. 2009, 304, doi:10.1016/j.mce.2009.02.021.

63. Hang, Y.; Stein, R. MafA and MafB activity in pancreatic β cells. Trends Endocrinol. Metab. 2011, 22.

64. Cano-Sancho, G.; Smith, A.; La Merrill, M.A. Triphenyl phosphate enhances adipogenic differentiation, glucose uptake and lipolysis via endocrine and noradrenergic mechanisms. Toxicol. Vitr. 2017, 40, 280–288, doi:10.1016/j.tiv.2017.01.021.

65. Green, A.J.; Graham, J.L.; Gonzalez, E.A.; La Frano, M.R.; Petropoulou, S.-S.E.; Park, J.-S.; Newman, J.W.; Stanhope, K.L.; Havel, P.J.; La Merrill, M.A. Perinatal triphenyl phosphate exposure accelerates type 2 diabetes onset and increases adipose accumulation in UCD-type 2 diabetes mellitus rats. Reprod. Toxicol. 2017, 68, 119–129, doi:10.1016/j.reprotox.2016.07.009.

66. Krumm, E.A.; Patel, V.J.; Tillery, T.S.; Yasrebi, A.; Shen, J.; Guo, G.L.; Marco, S.M.; Buckley, B.T.; Roepke, T.A. Organophosphate flame-retardants alter adult mouse homeostasis and gene expression in a sex-dependent manner potentially through interactions with era. Toxicol. Sci. 2018, 162, doi:10.1093/toxsci/kfx238.

67. Wang, D.; Yan, S.; Yan, J.; Teng, M.; Meng, Z.; Li, R.; Zhou, Z.; Zhu, W. Effects of triphenyl phosphate exposure during fetal development on obesity and metabolic dysfunctions in adult mice: Impaired lipid metabolism and intestinal dysbiosis. Environ. Pollut. 2019, 246, 630–638, doi:10.1016/j.envpol.2018.12.053.

68. Lee, Y.-M.; Ha, C.-M.; Kim, S.-A.; Thoudam, T.; Yoon, Y.-R.; Kim, D.-J.; Kim, H.-C.; Moon, H.-B.; Park, S.; Lee, I.-K.; et al. Low-Dose Persistent Organic Pollutants Impair Insulin Secretory Function of Pancreatic β-Cells: Human and In Vitro Evidence. Diabetes 2017, 66, 2669–2680, doi:10.2337/db17-0188.

69. Ward, A.B.; Dail, M.B.; Chambers, J.E. In vitro effect of DDE exposure on the regulation of B-TC-6 pancreatic beta cell insulin secretion: a potential role in beta cell dysfunction and type 2 diabetes mellitus. Toxicol. Mech. Methods 2021, 31, doi:10.1080/15376516.2021.1950251.

70. Santin, I.; Dos Santos, R.S.; Eizirik, D.L. Pancreatic beta cell survival and signaling pathways: Effects of type 1 diabetes-associated genetic variants. In Methods in Molecular Biology; 2016; Vol. 1433, pp. 21–54.

71. Nadal, A.; Soria, B. Glucose Metabolism Regulates Cytosolic Ca2+ in the Pancreatic β-Cell by Three Different Mechanisms. In Advances in Experimental Medicine and Biology; 1997; pp. 235–243.

72. Valdeolmillos, M.; Nadal, A.; Contreras, D.; Soria, B. The relationship between glucose-induced K+ATP channel closure and the rise in [Ca2+]i in single mouse pancreatic beta-cells. J. Physiol. 1992, 455, 173–186, doi:10.1113/jphysiol.1992.sp019295.

73. Martinez-Pinna, J.; Marroqui, L.; Hmadcha, A.; Lopez-Beas, J.; Soriano, S.; Villar-Pazos, S.; Alonso-Magdalena, P.; Dos Santos, R.S.; Quesada, I.; Martin, F.; et al. Oestrogen receptor β mediates the actions of bisphenol-A on ion channel expression in mouse pancreatic beta cells. Diabetologia 2019, 62, 1667–1680, doi:10.1007/s00125-019-4925-y.

74. Dos Santos, R.S.; Marroqui, L.; Velayos, T.; Olazagoitia-Garmendia, A.; Jauregi-Miguel, A.; Castellanos-Rubio, A.; Eizirik, D.L.; Castaño, L.; Santin, I. DEXI, a candidate gene for type 1 diabetes, modulates rat and human pancreatic beta cell inflammation via regulation of the type I IFN/STAT signalling pathway. Diabetologia 2019, 62, 459–472, doi:10.1007/s00125-018-4782-0.

75. Villar-Pazos, S.; Martinez-Pinna, J.; Castellano-Muñoz, M.; Alonso-Magdalena, P.; Marroqui, L.; Quesada, I.; Gustafsson, J.A.; Nadal, A. Molecular mechanisms involved in the non-monotonic effect of bisphenol-a on ca2+ entry in mouse pancreatic β-cells. Sci. Rep. 2017, 7, 11770, doi:10.1038/s41598-017-11995-3.

