## Supplemental Material for "Development of in vitro test methods in a model of human pancreatic β-cells to identify metabolism disrupting chemicals with diabetogenic activity"

**Dos Santos et al. - Supplementary Material**

**Table 1. List of primers used in this study.**

| Gene | Forward (5'-3') | Reverse (5'-3') |
|---|---|---|
| **Human β-actin** | CTGTACGCCAACACAGTGCT | GCTCAGGAGGAGCAATGATC |
| **Human *INS*** | GCTTCTTCTACACACCCAAGAC | CCACAATGCCACGCTTCT |
| **Human *PDX1*** | AAAGCTCACGCGTGGAAA | GCCGTGAGATGTACTTGTTGA |
| **Human *MAFA*** | TACAGGACGTGGACACCA | GTTCTCCGCTCAACCTCAG |
| **Human *GLUT2*** | TTCACTGCTGTCTCTGTATTCC | CTGACATGAAGATGGCACAAAC |
| **Human *GCK*** | GTCACCTGCAGCCTAATTACT | GCTTAGTGTCCTTCAGACAGATT |
| **Rat *Gapdh*** | AGTTCAACGGCACAGTCAAG | TACTCAGCACCAGCATCACC |
| **Rat *Ins1*** | ACCTTTGTGGTCCTCACCTG | AGCTCCAGTTGTGGCACTTG |
| **Rat *Ins2*** | TGTGGTTCTCACTTGGTGGA | CTCCAGTTGTGCCACTTGTG |
| **Rat *Pdx1*** | GGTATAGCCAGCGAGATGCT | TCAGTTGGGAGCCTGATTCT |
| **Rat *Mafa*** | AAGGAGGAGGTCATCCGACT | TCTGGAGCTGGCACTTCTCG |
| **Rat *Glut2*** | TCAGCCAGCCTGTGTATGCA | TCCACAAGCAGCACAGAGACA |
| **Rat *Gck*** | AGACTGACTATCCGGCTACAT | CCCAGAACTGTAAGCCACTC |

### Supplementary Figures

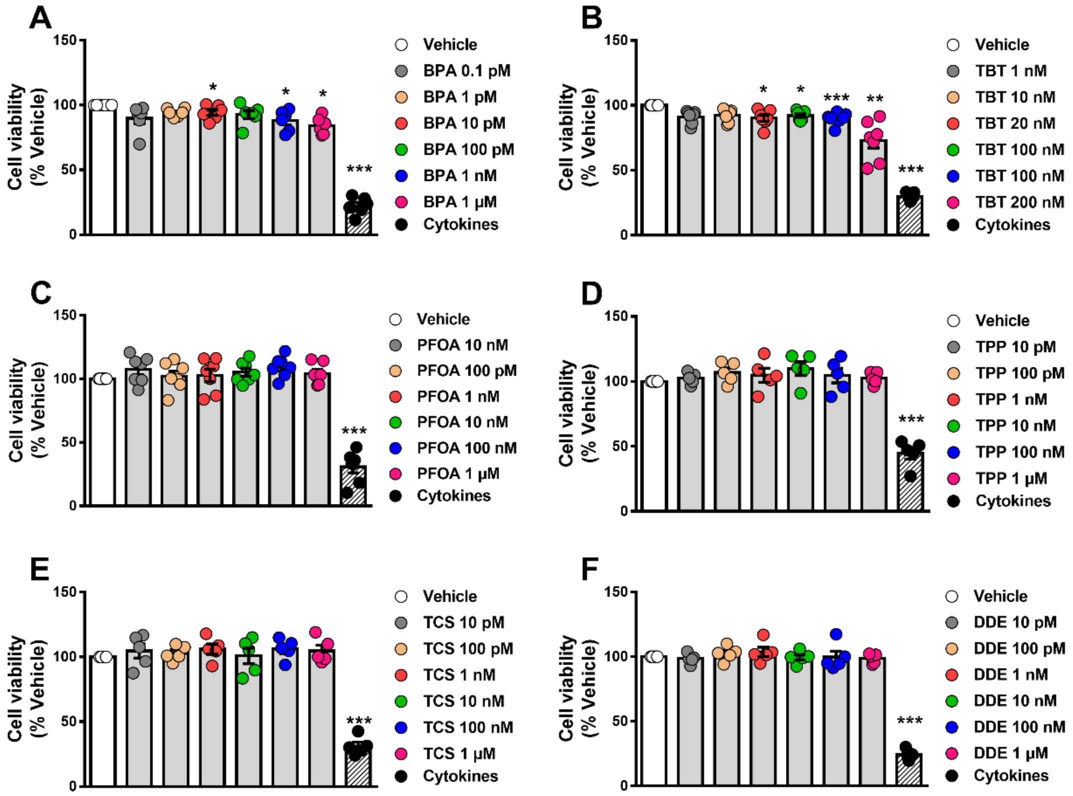


**Supplementary Figure 1. β-cell viability upon MDC exposure.** INS-1E cells were treated with vehicle (DMSO) or different doses of BPA (A), TBT (B), PFOA (C), TPP (D), TCS (E), or DDE (F) for 48 h. A cocktail of the cytokines IL-1β + IFNγ (10 and 100 U/ml, respectively) was used as a positive control. Cell viability was evaluated by MTT assay. Results are expressed as % vehicle-treated cells. Data are shown as means ± SEM of five to seven independent experiments. *p≤0.05, **p≤0.01 and ***p≤0.001 vs Vehicle. MDCs vs Vehicle by one-way ANOVA; Cytokines vs Vehicle by two-tailed Student's t test.

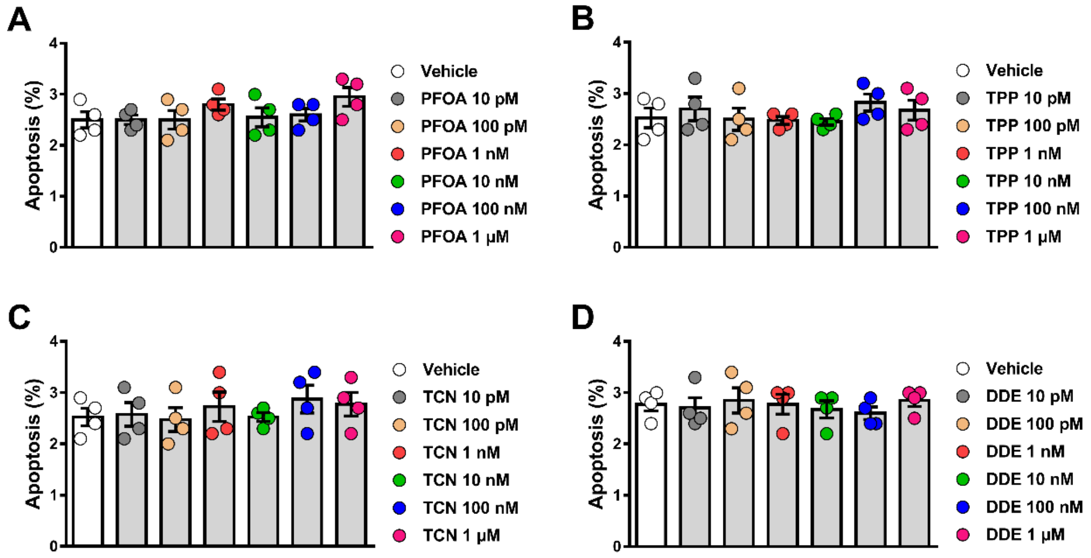


**Supplementary Figure 2. β-cell apoptosis upon MDC exposure.** INS-1E cells were treated with vehicle (DMSO) or different doses of PFOA (A), TPP (B), TCS (C), DDE (D) for 24 h. Apoptosis was evaluated using HO and PI staining. Data are shown as means ± SEM of four independent experiments.

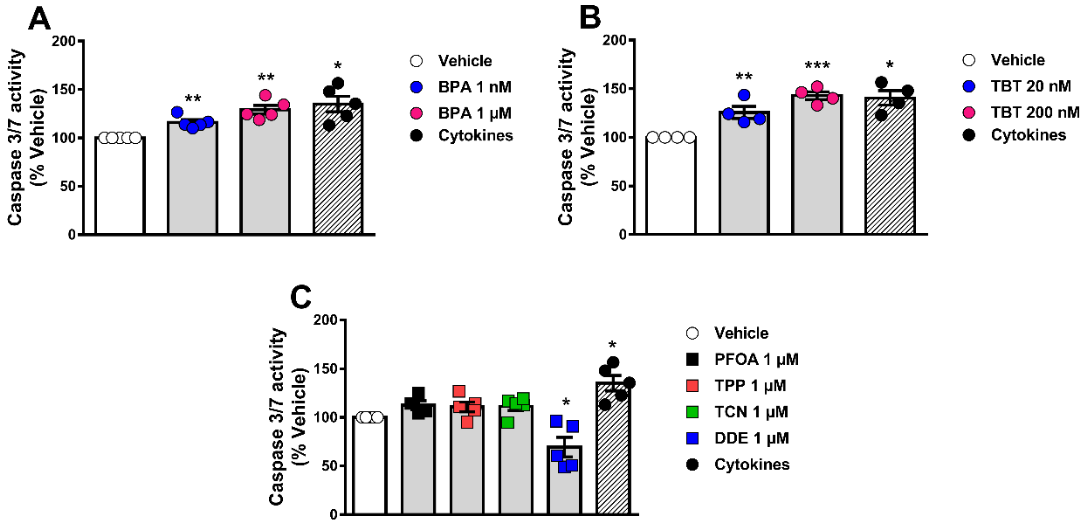


**Supplementary Figure 3. Caspase 3/7 activity upon MDC exposure.** EndoC-βH1 cells were treated with vehicle (DMSO) or different doses of BPA (A), TBT (B), PFOA, TPP, TCS, or DDE (C) for 48 h. A cocktail of the cytokines IL-1β + IFNγ (50 and 1000 U/ml, respectively) was used as a positive control. Caspase 3/7 activity was measured by a luminescent assay. Results are expressed as % vehicle-treated cells. Data are shown as means ± SEM of four to five independent experiments. *p≤0.05, **p≤0.01 and ***p≤0.001 vs Vehicle. MDCs vs Vehicle by one-way ANOVA; Cytokines vs Vehicle by two-tailed Student's t test.

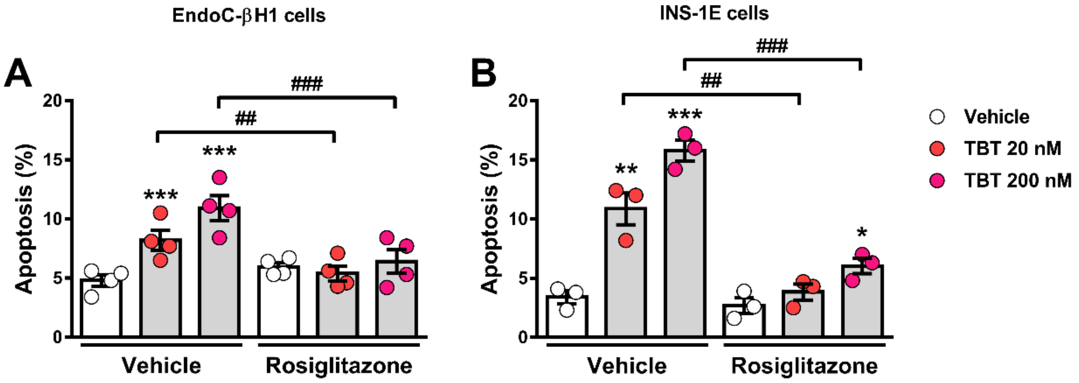


**Supplementary Figure 4. PPARγ is involved in TBT-induced β-cell apoptosis.** EndoC-βH1 (A) and INS-1E (B) cells were treated with vehicle (DMSO) or TBT (20 nM or 200 nM) in the absence or presence of 10 μM rosiglitazone for 24 h. Apoptosis was evaluated using HO and PI staining. Data are shown as means ± SEM of three to four independent experiments; *p≤0.05, **p≤0.01 and ***p≤0.001 vs its respective Vehicle; $^{\#\#}$p≤0.01, and $^{\#\#\#}$p≤0.001 as indicated by bars. Two-way ANOVA.

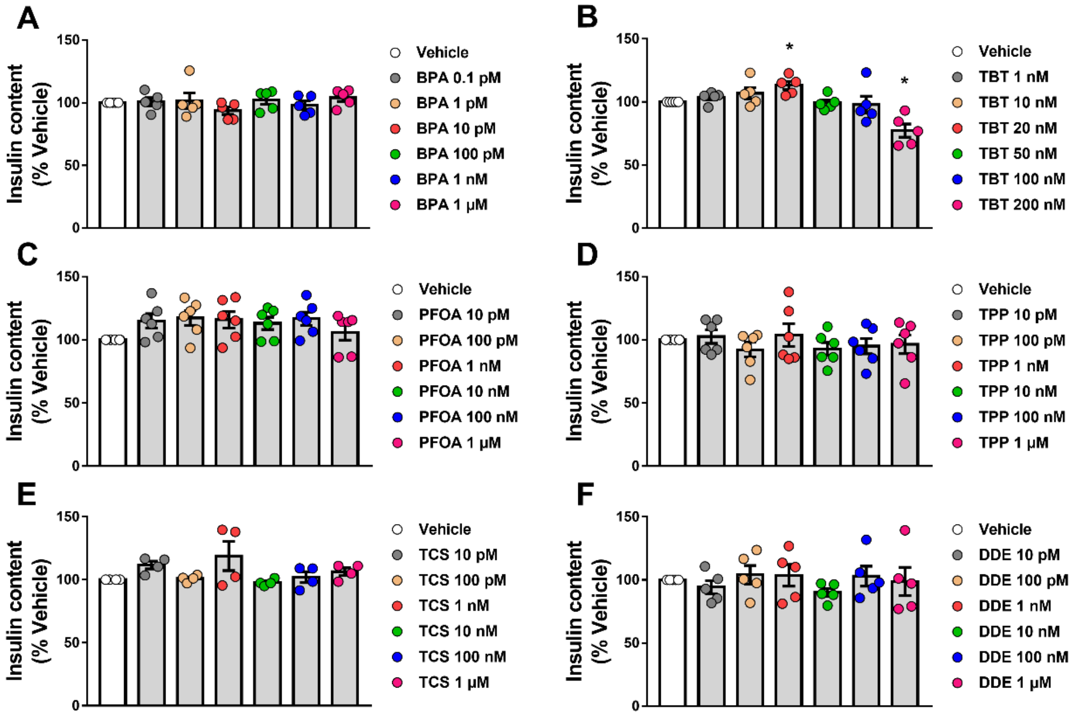


**Supplementary Figure 5. Insulin content upon MDC exposure**. EndoC-βH1 cells were treated with vehicle (DMSO) or different doses of BPA (A), TBT (B), PFOA (C), TPP (D), TCS (E), or DDE (F) for 48 h. Insulin content was measured by ELISA. Results are expressed as % vehicle-treated cells. Data are shown as means ± SEM of four to six independent experiments. *p≤0.05 vs its respective Vehicle. One-way ANOVA.

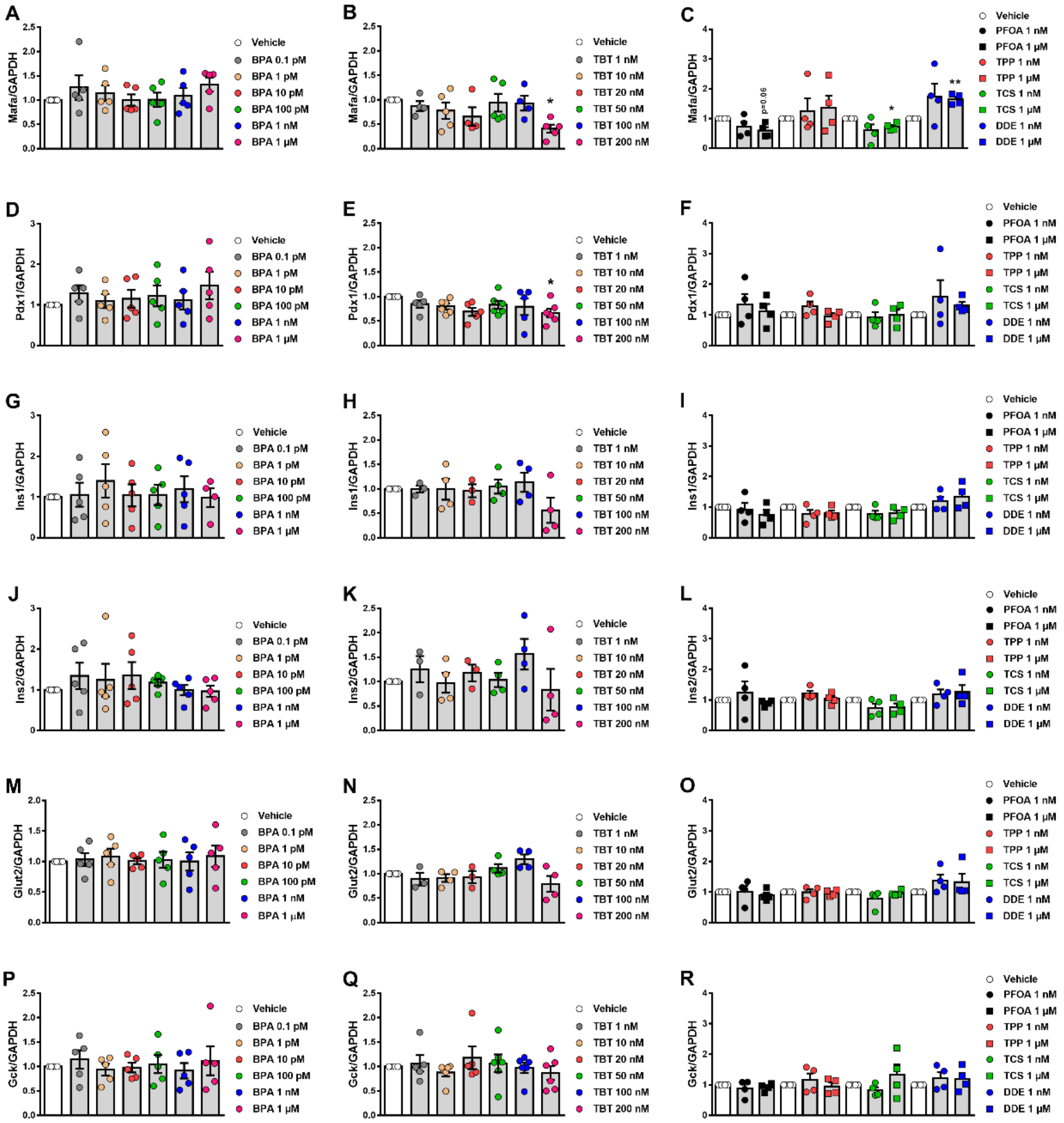


**Supplementary Figure 6. Gene expression upon MDC exposure**. mRNA expression of *Mafa* (A-C), *Pdx1* (D-F), *Ins1* (G-I), *Ins2* (J-L), *Glut2* (M-O), and *Gck* (P-R) was measured in INS-1E cells treated with vehicle (DMSO) or different doses of BPA (A, D, G, J, M and P), TBT (B, E, H, K, N and Q), PFOA, TPP, TCS, or DDE (C, F, I, L, O and R) for 24 h. mRNA expression was measured by qRT-PCR and normalized to the housekeeping gene *Gapdh*, and then by vehicle-treated cells (considered as 1). Data are shown as means ± SEM of three to six independent experiments. *p≤0.05 and **p≤0.01 vs its respective Vehicle. One-way ANOVA.
